# Structure of the bacterial cellulose ribbon and its assembly-guiding cytoskeleton by electron cryotomography

**DOI:** 10.1101/2020.04.16.045534

**Authors:** William J. Nicolas, Debnath Ghosal, Elitza I. Tocheva, Elliot M. Meyerowitz, Grant J. Jensen

**Affiliations:** Division of Biology and Biological Engineering, California Institute of Technology, Pasadena, California 91125; Howard Hughes Medical Institute, Pasadena, California 91125; Division of Medicine, Dentistry and Health Sciences, University of Melbourne Parkville VIC 3010, Australia; Department of Microbiology and Immunology, University of British Colombia, Vancouver, British Columbia V6T 1Z3

## Abstract

Cellulose is a widespread component of bacterial biofilms, where its properties of exceptional water retention, high tensile strength and stiffness prevents dehydration and mechanical disruption of the biofilm. Bacteria in the *Gluconacetobacter* genus secrete crystalline cellulose, with a structure very similar to that found in plant cell walls. How this higher-order structure is produced is poorly understood. We used cryo-electron tomography and focused ion beam milling of native bacterial biofilms to image cellulose-synthesizing *G. hansenii* and *G. xylinus* bacteria in a frozen-hydrated, near-native state. We confirm previous results suggesting that cellulose crystallization occurs serially following its secretion along one side of the cell, leading to a cellulose ribbon that can reach several microns in length and combine with ribbons from other cells to form a robust biofilm matrix. We were able to take direct measurements in a near-native state of the cellulose sheets. Our results also reveal a novel cytoskeletal structure, that we name the cortical belt, adjacent to the inner membrane and underlying the sites where cellulose is seen emerging from the cell. We find that this structure is not present in other cellulose-synthesizing bacterial species, *Agrobacterium tumefaciens* and *Escherichia coli* 1094, which do not produce organized cellulose ribbons. We therefore propose that the cortical belt holds the cellulose synthase complexes in a line, to form higher-order cellulose structures such as sheets and ribbons.

## Introduction

Humans rely on cellulose for building material, clothing and fuel^1–3^. More recently the polymer has sparked interest in the biotechnology field as a potential source of biofuel feedstock^4^, and in the biomedical industry as a biologically neutral scaffold to promote tissue regeneration^5,6^. Cellulose is a linear polymer of glucose molecules connected with *β*1-4 linkages by a glucosyltransferase. Individual linear glucan chains can pack via hydrogen bonding and Van Der Waals interactions in various ways to form different types of celluloses, with different properties^3,7,8^. The most common way glucan chains organize in nature is to form hydrogen-bonded planes which then stack into parallel layers via Van Der Waals interactions. These stacked layers give rise to cellulose I microfibrils, or “native cellulose”, that can then coalesce to form larger arrays. Because glucan chains pack in a regular lattice, cellulose I is considered crystalline. This form of cellulose is mainly found in plants, where it is a major structural element of the cell wall^9^.

In the prokaryotic world, cellulose is an important component of bacterial biofilms^10,11^, which increase cells’ tolerance for a range of biotic and abiotic stresses and enhance surface adhesion, cell cooperation and resource capture^10^. Cellulose-containing biofilms have also been implicated in pathogenicity, enabling bacteria to resist antibiotics and disinfection^12,13^. Most cellulose-synthesizing bacteria produce amorphous (non-crystalline) cellulose, but a few genera, including *Gluconacetobacter*, can produce cellulose I microfibrils like those found in plants. In *Gluconacetobacter*, these crystalline cellulose microfibrils can further aggregate into wide ribbon structures and larger arrays, giving rise to thick biofilms of pure cellulose I^14^.

Bacterial cellulose is synthesized by an envelope-spanning machinery called the Bacterial Cellulose Synthase (BCS) complex, encoded by the BCS operon and first identified in *Gluconacetobacter*^11^. While the components vary, most of the species encode BcsA, a component in the inner membrane that, with BcsB, catalyzes transfer of UDP-glucose to the nascent glucan chain^11,15,16^. BcsD forms a periplasmic ring that gathers glucan chains from several BcsA/B units^17^. BcsA and B are essential for cellulose synthesis *in vivo*, and BcsD is essential for the crystallization of nascent glucan chains^18^. BcsC forms a pore in the OM and very recent work has solved its crystallographic structure^19^. Consistent with previous data relying on sequence homology with the exopolysaccharide secretin components AlgE and AlgK from *P. aeruginosa*, BcsC is found to form an outer-membrane β-barrel pore at its C-terminal end, secreting the nascent elementary cellulose fibrils into the environment^18–22^. It is hypothesized that the elementary cellulose fibrils can aggregate with neighboring elementary fibrils upon secretion to form microfibrils^23,24^. Genes flanking the operon, *cmcax, ccpAX* and *bglxA*, are essential for cellulose crystallization and despite knowledge of their enzymatic functions, how they take part in this process is unclear^24–27^.

In the following report, the terms used to describe the cellulose assembly process are adapted from the ones defined in^24^, elaborating on the cell-directed hierarchical model for cellulose crystallization^7,28^. Glucan chains are linear polymers of β-1,4 linked glucose residues synthesized by a single catalytic site of a cellulose synthase. An elementary fibril (also termed mini-crystal in previous work^28–30^) is the product of the periplasmic aggregation of multiple glucan chains which is then extruded through a single BcsC subunit into the environment. Microfibrils result from the aggregation of several elementary fibrils, at least three according to earlier work^30^, outside the cell. These microfibrils can then crystallize into sheets that stack on each other to form ribbons.

Much work has already been done to understand the synthesis of crystalline cellulose^16,17,34–37,18,25–27,29,31–33^. In particular freeze-fracture electron microscopy (EM) studies have found that the *G. hansenii* BCS complexes are arrayed linearly along the side of the cell^29,36,37^, and this arrangement seems to determine the extracellular organization of cellulose I into ribbons^29,37^. How this linear arrangement is achieved is not known.

Here we used cryo-electron tomography (cryo-ET) of isolated cells and cryo-Focused Ion Beam (FIB)-milled biofilms to visualize native cellulose production in *G. hansenii* and *G. xylinus*, allowing the morphological characterization of the cellulose ribbons in a near-native state. We identified a novel cytoplasmic structure, which we call the cortical belt. We found that this cortical belt is absent from *Escherichia coli* 1094, which produces amorphous cellulose, and *Agrobacterium tumefaciens*, which produces crystalline microfibrils but not higher-order sheets, suggesting that the cortical belt functions to align BCS complexes to produce cellulose sheets.

## Results

### Cellulose is laid out in stacked sheets on one side of the cells

To visualize bacterial cellulose production, we used cryo-ET to image intact frozen-hydrated *G. hansenii* cells separated from their cellulose biofilm according to the original method from Brown et al. 1976. Previous work showed that newly synthesized cellulose ribbons are visible under the electron microscope at one hour post-separation^36^. To assure that the cells would have enough time to synthesize cellulose ribbons we imaged cells 5 hours (300 minutes) after separation. To confirm cellulose production, we stained cells with mitoTracker Deep Red FM to visualize membranes and Calcofluor-White to visualize cellulose. By confocal imaging, we observed cellulose filamentous structures tens of microns long (Fig. 1A and B, cyan arrowheads). We next plunge-froze cells at the same timepoint and imaged them by cryo-ET. The rod-shaped cells always lay flat on the grids, but their long axis was oriented randomly in the grid plane. Of 33 cells imaged, we found putative cellulose ribbons associated with 29 (88%), always on one side of the cell, including the top and bottom, and always aligned with the cell’s long axis (Fig 1C-E, yellow arrows). To confirm that the ribbon was in fact cellulose, we treated cells with cellulase and observed a large reduction in the occurrence of ribbons in cryo-EM and negative stained images (Supplemental figure 1, yellow arrowheads). Instead, we observed aggregated material we think is likely partially digested cellulose (Supplemental figure 1F, orange arrowheads).

**Figure 1.**
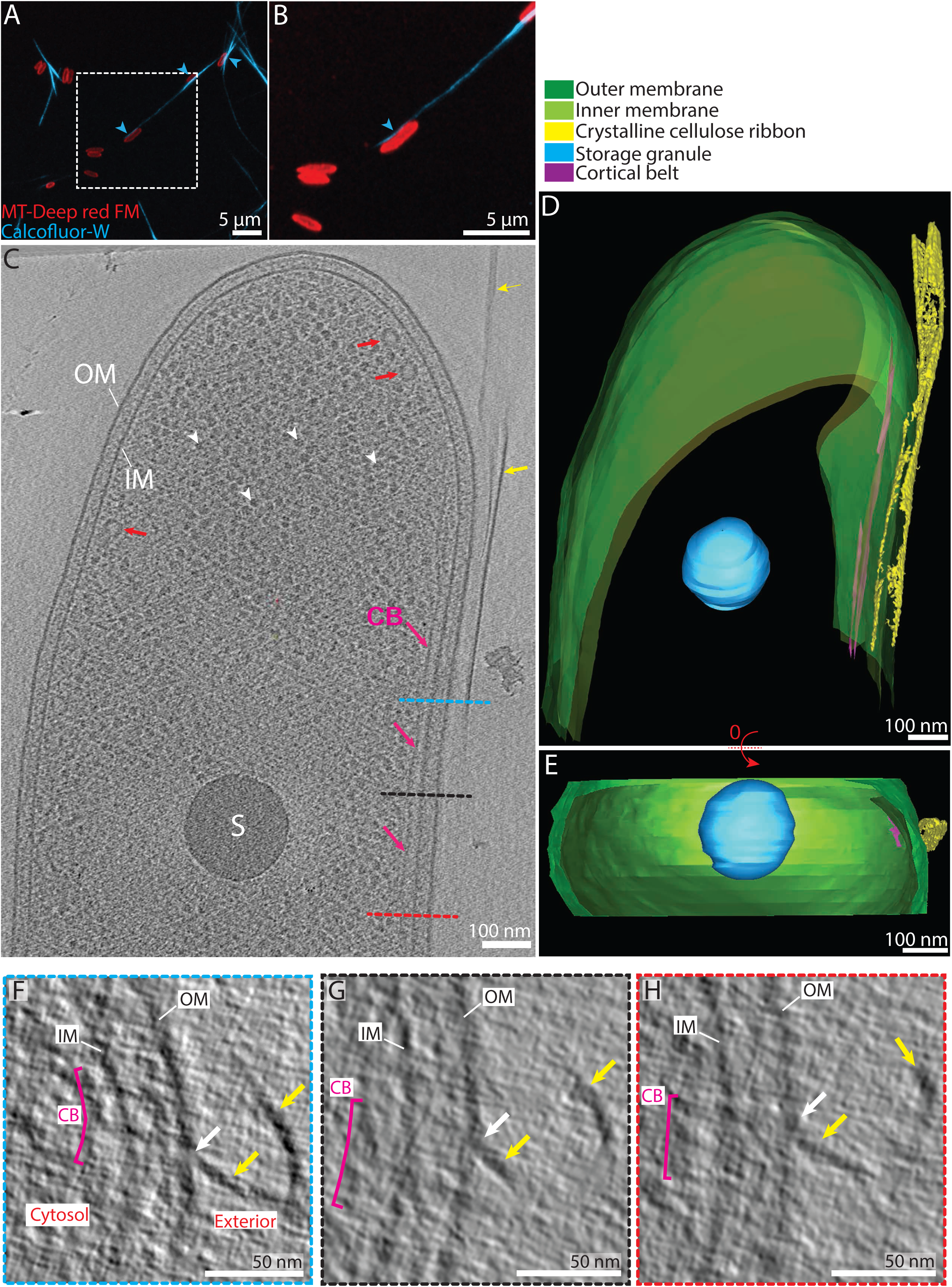
Interactions between the bacterial envelope and the cellulose ribbon: the “tight” configuration. (**A**) Confocal-Airy scan optical slices show representative examples of *G. hansenii* cells in red (MitoTracker Deep Red FM) displaying the cellulose ribbon on their side in cyan (Calcofluor-white). (**B**) Enlarged view indicated by white dashed rectangle in (A). The cellulose structure is clearly seen closely appended to one side of the cell (cyan arrowheads). (**C**) 9-nm thick tomographic slice showing the typical *G. hansenii* cell harboring the cellulose ribbon on its right side (yellow arrows). White arrowheads point to ribosomes and red arrows point to cytosolic vesicles. Here and below, IM: Inner-membrane; OM: Outer-membrane; S: Storage granule; CB: Cortical belt. (**D**) Manual segmentation of the cell shown in (C). (**E**) Rotated segmented volume shown in (D) showing the very close contact between the cellulose ribbon (yellow) and the outer membrane (green). (**F-H**) Transverse 9-nm thick tomographic slices through the bacterial envelope of the cell shown in (C) at the levels indicated by the blue, black and red dashed lines, respectively. Two cellulose sheets (yellow arrows) are seen. One interacts with the OM all along (white arrow). Our interpretation is that integration of the cellulose fibers into the sheet occurs immediately upon secretion.

The spatial relation between the cellulose ribbons and the OM was examined. In 3 out of the 29 tomograms, the cellulose ribbon was observed running beneath or on top of the cell, causing it to be normal to the electron beam thus inherently not well resolved and difficult to assess its spatial relation with the OM. Therefore, data from these 3 tomograms was excluded for these measurements. In the remaining tomograms two distinct configurations were observed: a “tight” configuration in 23 out of 26 tomograms (88%), where the average outer membrane (OM)-to-ribbon distance was 16 ± 5 nm (n = 23) (Fig. 1C-H, supplemental video 1), and a “loose” configuration in 3 out of 26 tomograms (12%), where the average OM-to-closest sheet distance was 99 ± 49 nm (n = 3) (Fig. 2). Among the tomograms showing a “tight” configuration, 17 out of 23 (65%) displayed multiple clear direct contacts between the OM and the ribbon (Fig. 1F-H, white arrows). Tomograms in the “loose” configuration exhibited ribbons that seemed detached from the OM, with an increased OM-to-closest sheet distance compared to the “tight” configuration (Fig. 2E). All three tomograms presented disorganized aggregates bearing a mesh-like appearance between the OM and the ribbon (Fig. 2A-D, orange asterisks and dashed bracket). These aggregates always connected to the ribbon (Fig. 2A, black lined orange arrows). Similar cellulose aggregates have been seen previously by negative staining^23^.

**Figure 2.**
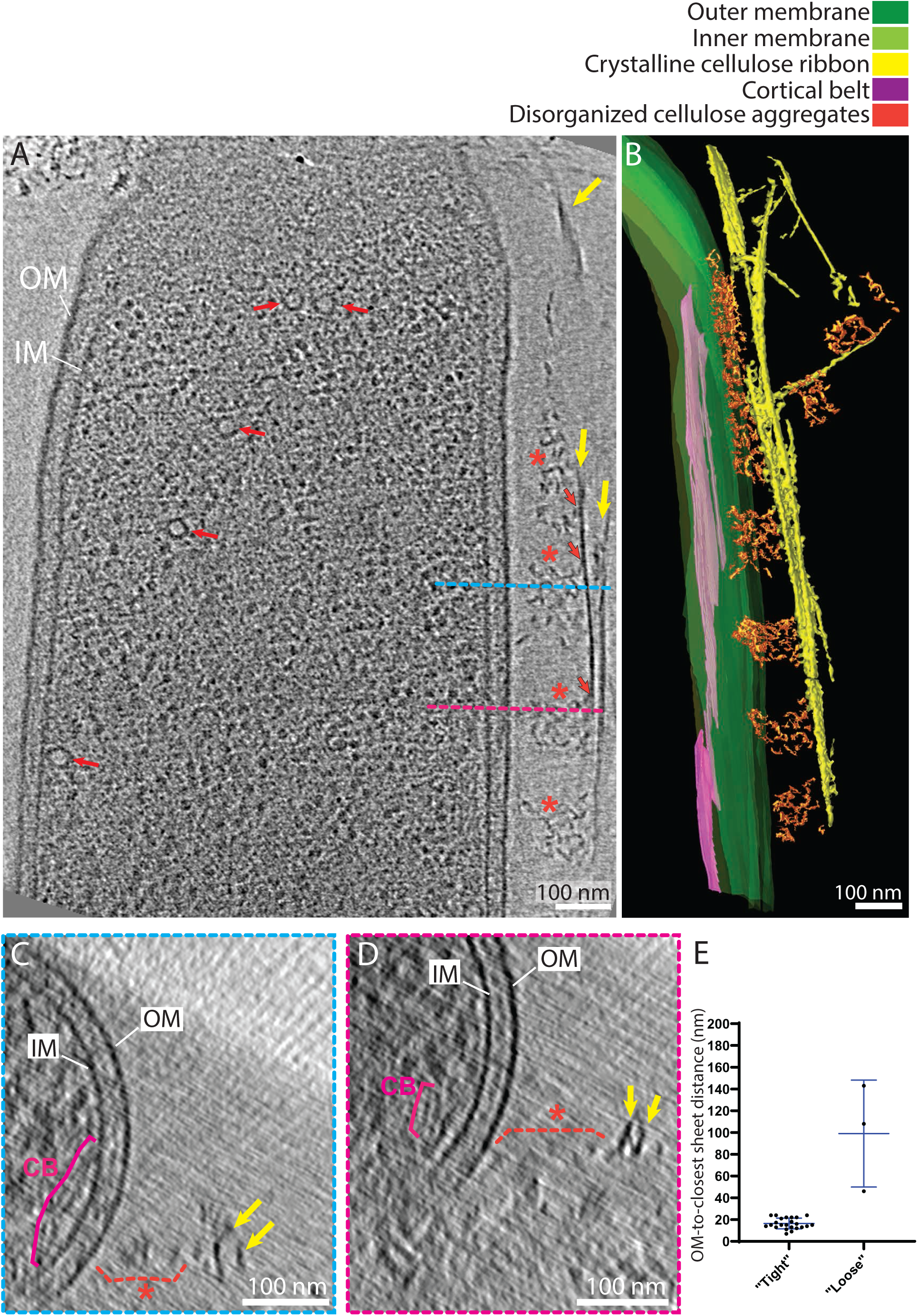
Interactions between the bacterial envelope and the cellulose ribbon: the “loose” configuration. **(A)** 9-nm thick tomographic slice showing a cell where aggregates of disorganized cellulose (orange asterisks) occur between the ribbon (yellow arrows) and the OM. Note the cortical belt (CB) cannot be seen in this slice. Black line orange arrows indicate points of contact between the cellulose sheet and the disorganized aggregates. Red arrows point to vesicles. (**B**) Manual segmentation of the tomogram in (A) showing these disorganized aggregates in 3-D. (**C-D**) Transverse 9-nm thick tomographic slices through the envelope of the cell shown in (A) at the levels indicated by the blue and pink dashed lines highlighting the distance between the two cellulose sheets (yellow arrows) and the OM and the presence of the disorganized clusters (orange dashed brackets). (**E**) Plot showing the OM-to-closest sheet distance in the two types of configuration.

These cells and their cellulose structures (the ribbons) were imaged in a near-native (frozen hydrated) state, allowing measurement of their native dimensions. In our description of the cellulose ribbons below, by length we mean the dimension parallel to the long axis of the cell (Fig. 3A). By thickness we refer to the dimension normal to the cell surface (Fig. 3A, black inset). By width we refer to the dimension tangential to the cell surface (Fig. 3B). The cellulose ribbons we observed were very similar to what has been seen previously by negative stain EM^23,36^. Ribbons comprised long flexible sheets, too long to be measured by cryo-ET because they are never entirely in the field of view. All structures in an electron tomogram suffer from the missing wedge artefact, inherent to electron tomography, which causes elongation in the direction parallel to the electron beam^38,39^. For this reason, measurements of the width of the cellulose sheets are systematically overestimated. Sheet width was estimated at 38 ± 14 nm (n = 45) (Fig. 3C). These sheets then stack into a ribbon (2.3 ± 0.9 sheets on average; n = 24), with a variable inter-sheet distance (16 ± 7 nm; n = 23). Inter-sheet distance was accurately measured peak-to-peak (Fig. 3D), which encompasses 2 halves of the two neighboring sheets’ density and the space between them (Fig. 3A, black inset). Because the apparent thickness of single densities in cryo-ET is strongly affected by the defocus applied, individual cellulose sheet thickness measurements will be overestimated. Therefore, we can only say confidently that they are thinner than the inter-sheet distance. Despite careful inspection, although densities could be seen in the periplasmic space, we did not recognize a consistent shape which we could confidently attribute to the BCS machinery. This is likely due to the large cell diameter (∼800nm), and the small size and/or flexibility of the BCS complexes.

**Figure 3.**
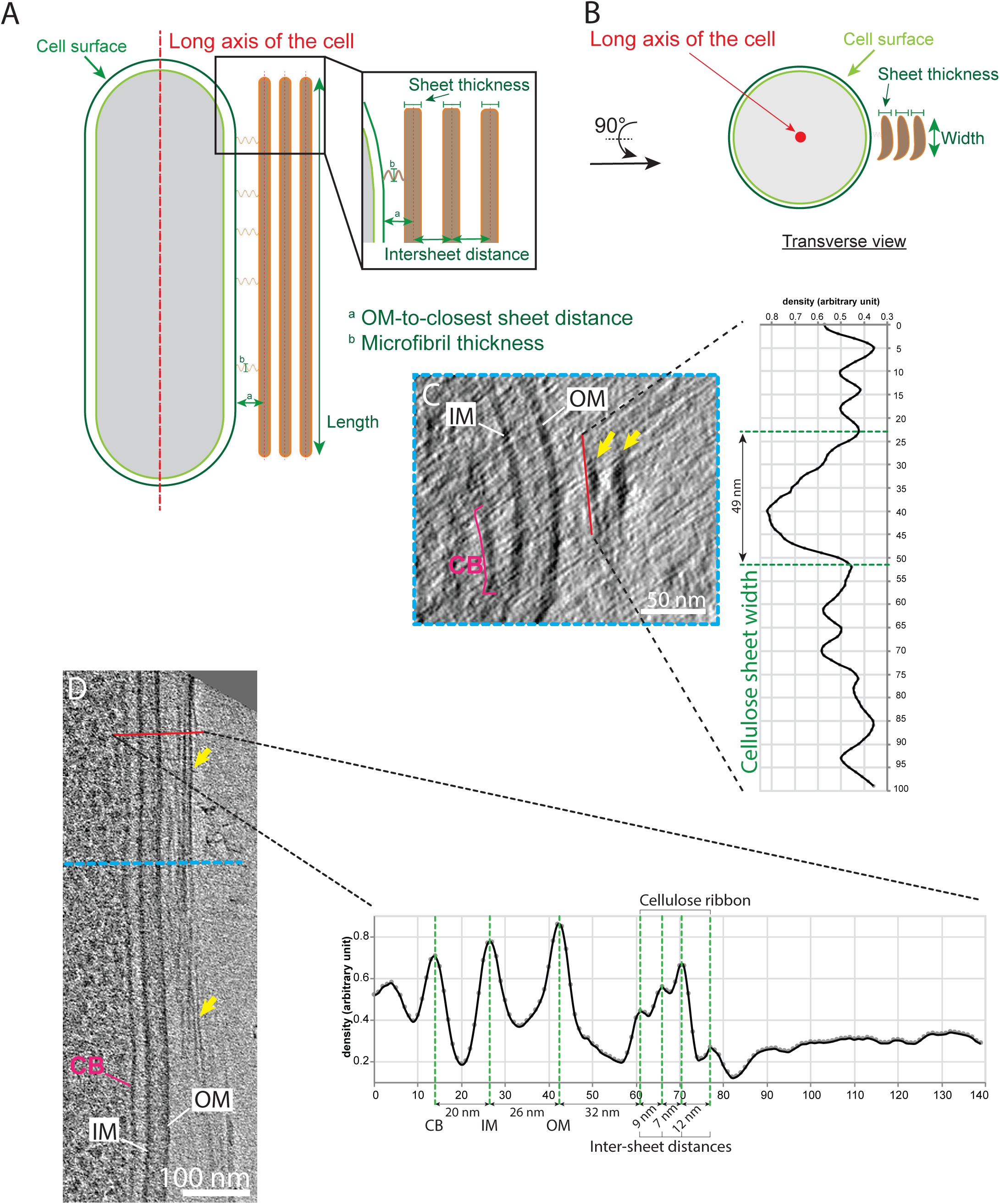
Cellulose sheet dimensions. (**A-B**) Longitudinal and transverse schematic depiction defining the different dimensions measured, namely OM-to-sheet distance, sheet width and inter-sheet distance. Identical terminology is used for the measurements of the cortical belt. (**C**) Transverse 12-nm thick slice of the bacterial envelope of the cell shown in (D) at the level indicated by the blue dashed line. The yellow arrows highlight the two stacked sheets. On the right, the average density profile along the red line demonstrates how the cellulose sheet widths were estimated. Vertical axis is length in nm along the red line and horizontal axis is the normalized electron density. (**D**) 12-nm thick tomographic slice showing the typical organization of the bacterial envelope on the side where cellulose sheets (yellow arrows) are being synthesized. The average density profile on the right taken along the red line shows the CB-IM, IM-OM OM-sheet and inter-sheet distances (green dashed lines).

### Sheets arise from the stacking of microfibrils

To visualize earlier stages of cellulose synthesis, we plunge-froze cells at earlier timepoints after separation from the biofilm. A total of 6 and 15 tomograms were acquired at 13- and 20-minutes post-separation, respectively. At 13 minutes (the most quickly we could complete plunge freezing), no cells exhibited a cellulose ribbon, however, disorganized aggregates were observed in the vicinity of 1 out of the 6 tomograms. At 20 minutes post-separation, cellulose ribbons were observed adjacent to the cell in 9 out of 15 tomograms (64% versus 88% (n = 33) at 300 minutes post-separation) (Fig. 4A). Out of these 9 cells harboring an adjacent cellulose ribbon, 3 had it on the top or bottom of the cell and were excluded from the analysis for the same reason explained above. Therefore, the analysis of the OM-ribbon interface was conducted on the remaining 6. The cellulose ribbons observed at 20min post-separation comprised only one cellulose sheet (n = 6) (Fig. 4B). Four out of these 6 tomograms (67%) exhibited a “tight” configuration. The average OM-to-closest sheet distance of 14 ± 3 nm (n = 4) was not significantly different from the 300 minutes post-separation “tight” configuration average OM-to-closest sheet distance (Fig. 4C-D). The two other tomograms bore ribbons in the “loose” configuration, i.e. at an OM-to-closest sheet distance >40 nm with disorganized aggregates in-between. These “loose” ribbons had an OM-to-closest sheet distance of 43 and 59 nm, respectively. The disorganized aggregates visible at 20 minutes post-separation emanated perpendicularly from the OM to connect to the nascent cellulose sheet. They were thinner than the ones observed at 300 minutes post-separation and rod-shaped (Fig. 4E-F, red arrowheads). Average density profiles normal to the direction of the cylindrical-shaped densities were traced to estimate their diameter (Fig. 4G). We again emphasize the inherent overestimation of such measurements due to defocus. The average estimates on the two cells, 11 ± 2 nm (n = 12) and 6.5 ± 1 nm (n = 4), respectively (Fig. 4G), therefore establish upper limits of the true diameter. These estimates are also less than the above-measured inter-sheet distances (Fig. 4H). Because elementary fibrils are thought to be between 1.5 and 6 nm in thickness^14,36,37^, we interpret these structures as microfibrils composed of several elementary fibrils. The variability of the microfibril diameter measurements between cells (Fig. 4G, Cells #1 and #2) suggests these structures can contain a varying number of elementary fibrils more-or-less tightly packed together. This configuration is reminiscent of what was seen in previous studies of microfibrils coming out of clusters of pores^23,36^ and likely represents an early stage of cellulose sheet formation that has been mechanically disturbed. Sheets at 20 minutes post-separation had an estimated width of 25 ± 8 nm (n = 6) (Fig. 4I), smaller than those at 300 minutes.

**Figure 4.**
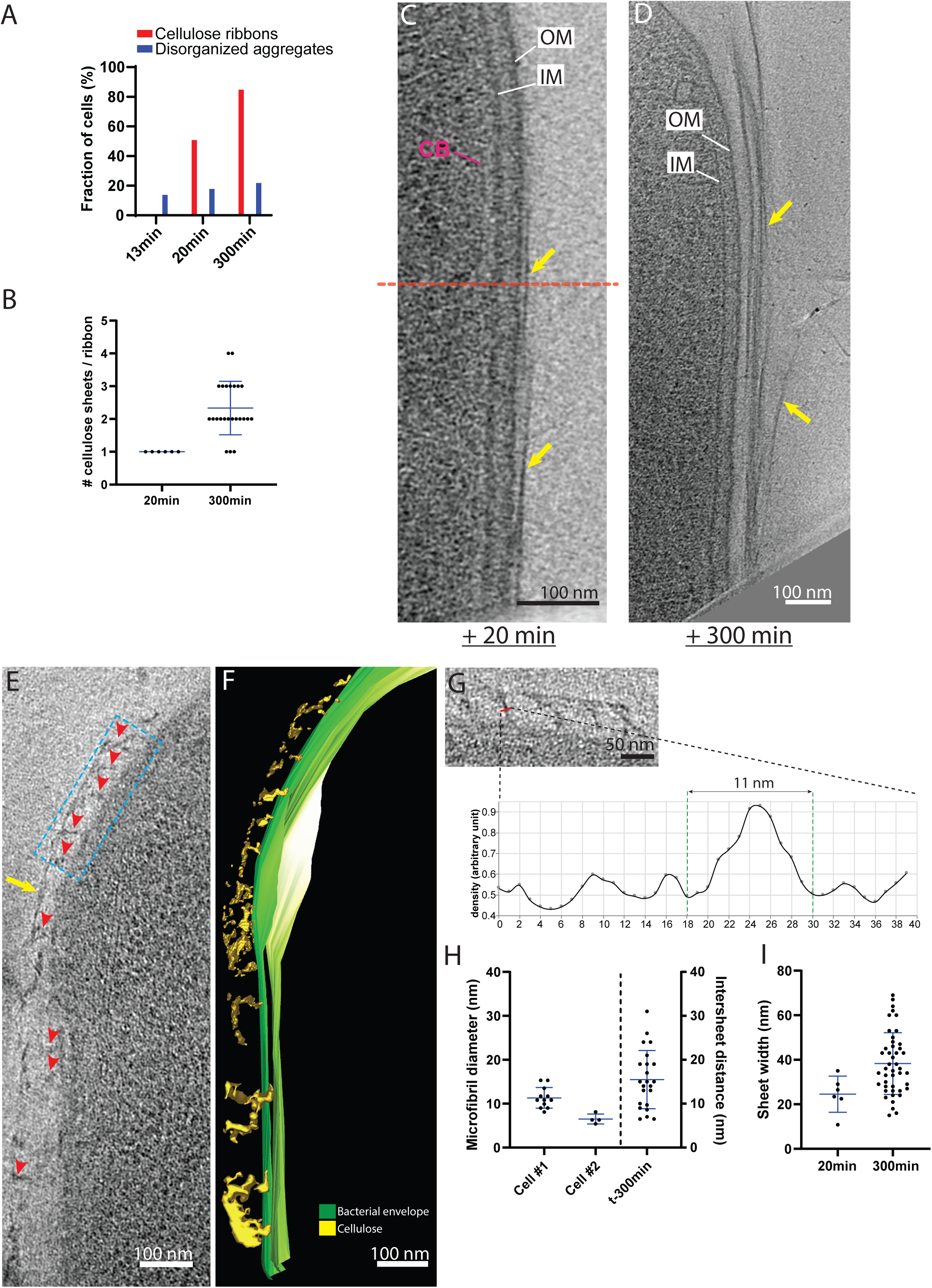
The cellulose ribbon is a composite structure made of stacked sheets. (**A**) Percentages of cells exhibiting disorganized aggregates (blue) and cellulose ribbons (red) at 13-, 20- and 300minutes post-separation. While disorganized aggregate occurrence is steady, there is an increase in the occurrence of cellulose ribbons over time. (**B**) Number of cellulose sheets composing the ribbons as a function of time after cell separation. (**C**) Composite image composed of 10-nm thick tomographic slices spaced by 24 nm in Z, of a cell 20 minutes post-separation in the “tight” configuration. The cellulose ribbon is thin (yellow arrows), composed of one sheet immediately adjacent to the OM. Limits of the two original images are indicated by the red dashed line. (**D)** 11 nm thick tomographic slice of a cell 300 minutes post-separation. The cellulose ribbon (yellow arrows) is large and composed of multiple sheets. (**E**) Nascent cellulose sheet 20 minutes post-separation (yellow arrow). Putative microfibrils can be seen coming out perpendicularly from the outer membrane (red arrowheads). (**F**) Corresponding manual segmentation of (E). (**G**) Enlarged view of the blue boxed region in (E). Below is the average density profile showing the estimation of the diameter of one putative microfibril (red line). (**H**) Estimated diameters of microfibrils observed at 20-minutes post-separation in the two cells where they are visible (left vertical axis) as in (E) and the inter-sheet distances measured in the 300-minutes post-separation cellulose ribbons (right vertical axis). (**I**) Sheet width estimations at 20- and 300-minutes post separation.

These results show that 1) the microfibrils emanating from the OM have roughly the same thickness as the cellulose sheet, 2) sheet width seems to increase over time and 3) the number of cellulose sheets comprising a ribbon increases over time.

### A novel cytoplasmic structure is associated with cellulose production

We next examined the interior of *G. hansenii* cells during cellulose synthesis. These cells had extensive cytoplasmic vesicles in the center and at the periphery of the cell (Supplemental figure 2). The most notable feature we observed was another ribbon-like structure closely associated with the inner membrane (24 ± 4 nm from it; n = 19, for an example peak-to-peak measurement see Fig. 3D) and several hundred nanometers in length (Fig. 5A, purple arrows). We found it in 90% of cells with a cellulose ribbon (n = 29), always on the same side as, and underlying, the nascent cellulose sheet (Fig. 5B-C, supplemental video 2). This cytoplasmic structure is not a tube but rather a stack of sheet-like structures, 47 ± 23 nm wide (n = 10), parallel to the inner membrane and spaced (peak-to-peak) by 15 ± 5 nm (n = 7) (Fig. 5D-F). We refer to it here as the “cortical belt”.

**Figure 5.**
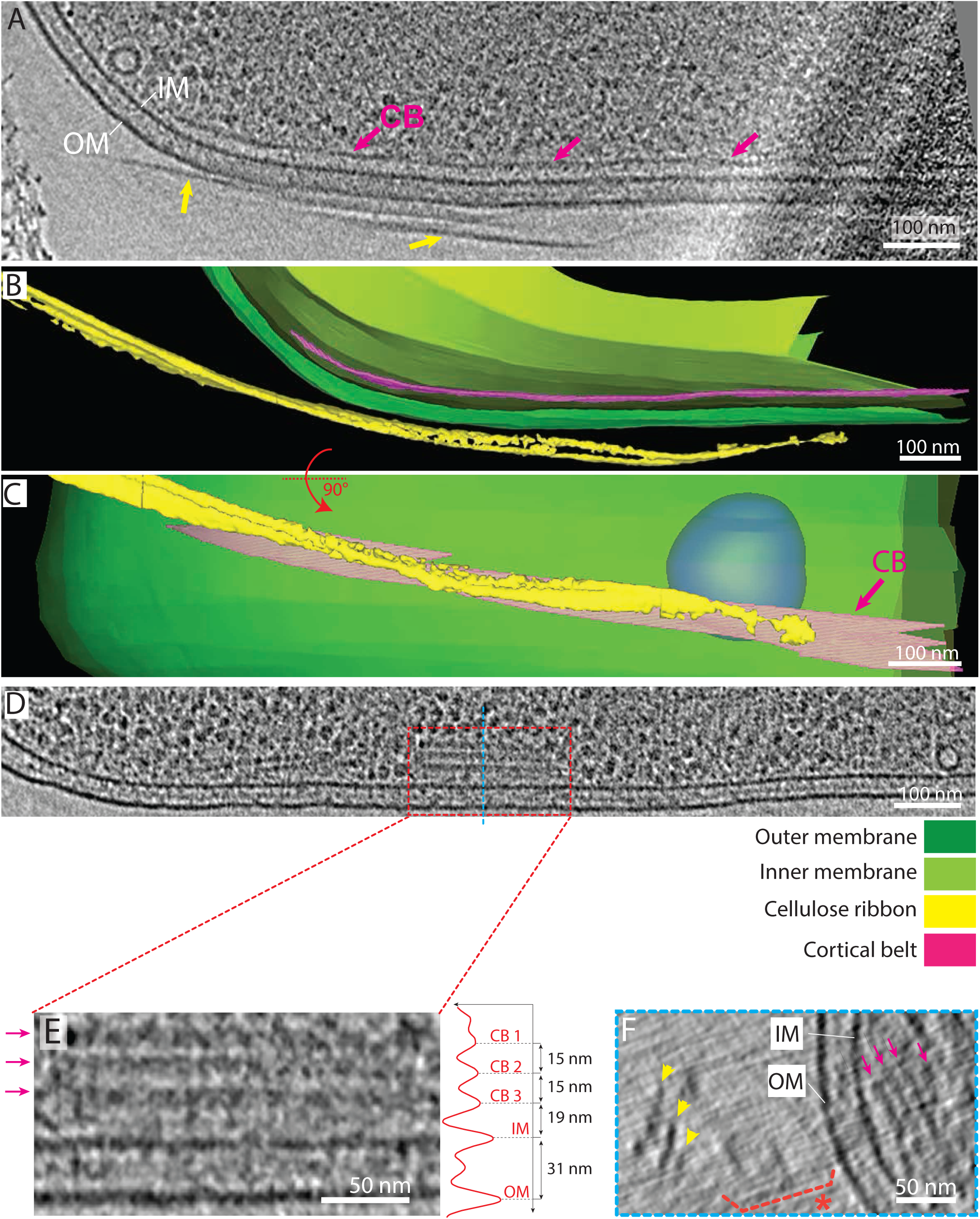
The cortical belt lies below the cellulose ribbon in the cytoplasm. (**A**) 9-nm thick tomographic slice showing a representative cortical belt (purple arrows) just inside the IM and proximal to the cellulose ribbon on the outside of the cell (yellow arrows). (**B**) Manual segmentation of the tomogram shown in (A) highlighting the cellulose ribbon and the cortical belt. (**C**) Same segmentation rotated 90° about the long axis of the cell shows how the cortical belt and the cellulose ribbon follow the same trajectory. (**D**) 9-nm thick tomographic slice taken from the same tomogram as in figure 2, showing one out of several cases where the cortical belt presented stacked layers (red dashed box). (**E**) Enlarged view of the red dashed boxed region in (D) showing the arrangement of the stacked layers. On the right is a density profile displayed normal to the cortical belt to measure the inter-layer distance (15 nm). (**F**) Transverse 9-nm thick tomographic slice of the cell region shown in (D), at the level indicated by the blue dashed line, highlighting stacked layers of the cortical belt. The cellulose ribbon can be seen at a distance (yellow arrowheads) with disorganized aggregates in between (orange dashed brackets and asterisk).

### Structural hallmarks of crystalline cellulose synthesis are also present in intact biofilms

It is possible that separating bacteria from the cellulose mat for whole cell cryo-ET imaging could have altered structures associated with cellulose synthesis. We therefore imaged *G. hansenii* cells *in situ* in young cellulose biofilms grown on gold grids. We imaged biofilms after 3 or 6 hours before plunge-freezing in hope of visualizing any change in the ordering of the fibers or the aspect of the cells over the course of biofilm growth. To access cells within the 5- to 10-micron thick biofilm, we used cryo-FIB milling to generate thin (∼200 nm) lamellae suitable for imaging by cryo-ET (Fig. 6A-C). In a total of 19 analyzed tomograms (9 and 10 tomograms for 6h and 3h biofilms, respectively, Table 1), we observed fields of living and dead bacteria encased in a matrix of bundled cellulose ribbons at both time points (Fig. 6D-E and supplemental video 3). Overview tomograms (low magnification with low total dose) and high-resolution composite images of the lamellae allowed extraction of positional information of the cells in relation to the biofilm. There were 0.10 ± 0.02 and 0.27 ± 0.04 cells/um^2^ and 15% and 28% of the volume of the lamellae was occupied by cells at 3 and 6h time points, respectively (Fig. 6F) (n = 6 and 4 lamellae, respectively). This approximate 2-fold increase in cell density from a 3-hour to a 6-hour biofilm suggests that cell division is occurring during biofilm growth. As dead cells can be easily identified by their appearance (Fig. 6D, red asterisks), the live-to-dead cell ratio was calculated at 0.9 ± 0.1 in both 3- and 6-hour biofilms, revealing no increase in the proportion of dead cells between these two timepoints (Fig. 6G). Because lamellae give access to the native organization and layering of the cells within the biofilm, the depth of dead/living cells within the biofilm was assessed by measuring their distance from the leading edge of the lamella (see methods). No trend between cell depth within the biofilm and state of the cells was detected (Fig. 6H).

| Whole cell tomography |  |  |  |  |  |
| --- | --- | --- | --- | --- | --- |
| Species | Condition/strain | Lamellae | Tomograms | VPP | Source |
| *G. hansenii* (ATCC 23769) | Untreated - 13min post sep | N/A | 6 | - | ATCC |
|  | Untreated - 20min post sep | N/A | 14 | - |  |
|  | Untreated - 300min post sep | N/A | 33 | 24 |  |
|  | Cellulase 0.2g/L | N/A | 4 | - |  |
| *G. xylinus* (ATCC 700178) | Untreated - 300min post sep | N/A | 8 | - | ATCC |
| *A. tumefaciens* C58<sup>1,3</sup> | WT<sup>2</sup> | N/A | 47 | - | gift from Patricia Zambrisky (UC Berkeley) to Elitza Tocheva |
|  | A139<sup>2</sup> | N/A | 10 | - | gift from Patricia Zambrisky (UC Berkeley) to Elitza Tocheva |
|  | AD348<sup>2</sup> | N/A | 1 | - | gift from Anath Das (University of Minnesota) to Debnath Ghosal |
|  | AD1484<sup>2</sup> | N/A | 1 | - | gift from Anath Das (University of Minnesota) to Debnath Ghosal |
|  | JX148<sup>2</sup> | N/A | 4 | - | gift from Patricia Zambrisky (UC Berkeley) to Elitza Tocheva |
|  | NT1<sup>2</sup> | N/A | 2 | - | gift from Patricia Zambrisky (UC Berkeley) to Elitza Tocheva |
| Tomography on milled lamellae |  |  |  |  |  |
| Species | Condition/strain | Lamellae | Tomograms |  | Source |
| *G. hansenii* (ATCC 23769) | Native biofilm - untreated | 12 | 33 | 3 | ATCC |
| *E. coli* 1094 induced for cellulose synthesis | Untreated | 2 | 6 | - | gift from Jean Marc Ghigo (Institut Pasteur) |
|  | Cellulase 0.2g/L | 1 | 2 | - |  |

<sup>1</sup> C58 or ATCC 33970 is the wild type *A. tumefaciens*
<sup>2</sup> All these strains have a C58 background.
<sup>3</sup> These strains were imaged for other purposes, but used here as well

**Figure 6.**
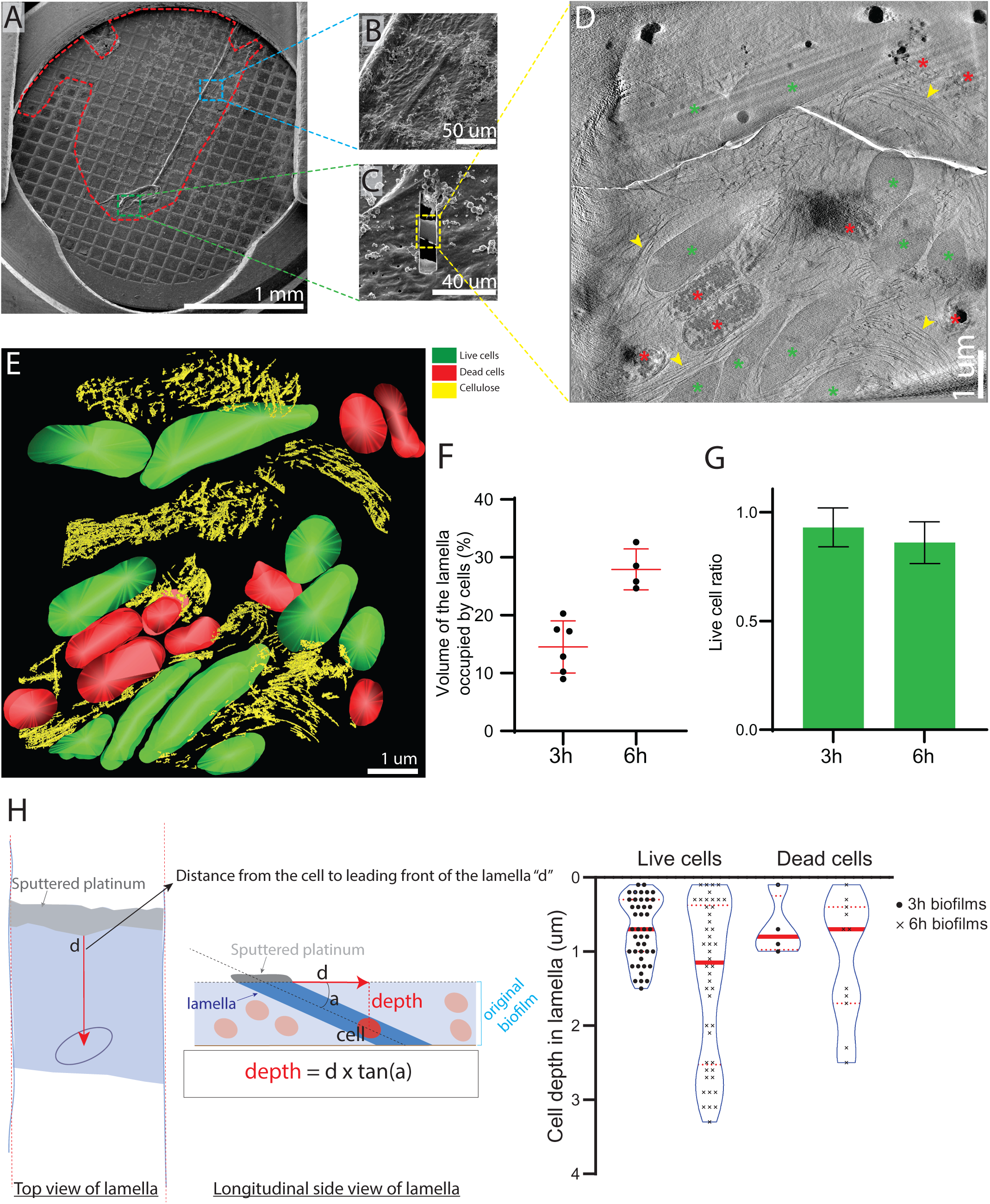
FIB-milling through native *G. hansenii* biofilms. (**A**) cryo-SEM overview of a 6-hour biofilm (outlined in red) grown on a gold quantifoil grid. (**B**) cryo-SEM view of a thick biofilm area (boxed in blue in (A)). Wrinkles in the biofilm are typical of a biofilm a few microns thick. (**C**) Milled lamella (boxed in yellow) from the green boxed region shown in (A). (**D**) 23-nm thick tomographic slice of a low mag tomogram taken on the lamella shown in (C). Living (when frozen) and dead cells are visible (green and red asterisks, respectively) and large cellulose arrays can be seen filling the gaps between the cells (yellow arrowheads). (**E**) Manual segmentation of the tomogram shown in (D). (**F**) Fraction of the lamella volume occupied by the cells was assessed for each lamella (**G**) Live cell ratio in 3h and 6h biofilms. (**H**) Violin boxplots reporting the absolute depth of the live and dead cells within the biofilms grown for 3 and 6 hours. The dashed red lines indicate the first and third quartiles and solid red line represents median. This shows that while the biofilms get thicker with time, the ratio of live-to-dead cells appears constant through depth and time. Method of calculation is detailed on the left of the panel and in the methods section. Lamella is drawn in blue, with the platinum coated leading edge represented in gray.

In all 19 tomograms (combining 3h and 6h lamellae), we observed numerous cellulose ribbons surrounding the cells (Fig. 7A, yellow arrowheads). In 5 out of the 19 tomograms (26%), a cellulose ribbon was closely appended to the cell’s OM, as we previously had seen in separated cells (Fig. 7B-C, dark-lined yellow arrowhead). Among those 5 tomograms, 4 showed a cortical belt adjacent to the cellulose ribbon (Fig. 7B-D and supplemental video 3). The OM-to-cellulose ribbon distance (19.2 ± 8 nm, n = 4) and inner membrane to cortical belt distance (22 ± 2 nm, n = 4) were very similar to those measured before in separated cells. In 5 out of the 10 tomograms in 3h biofilm lamellae, disorganized cellulose aggregates were observed connected to well-ordered ribbons just as in the separated cells, whereas this was never observed in the 6h biofilms. This suggests that crystallization is disrupted more often in early biofilm growth (Fig. 7E-G, orange dashed lining). Because *Gluconacetobacter* cells are thick, electron transmittance in the central region of the cytoplasm is very low when imaging whole cells, making it difficult to visualize this area. Reducing sample thickness to approximately 200 nm by cryo-FIB-milling allowed us to observe these central regions with greater contrast and visualize the extensive vesicle network deep inside the cell (Fig. 7E, white arrowheads). Overall, the morphology of cells and cellulose structures in a biofilm was the same as in isolated cells.

**Figure 7.**
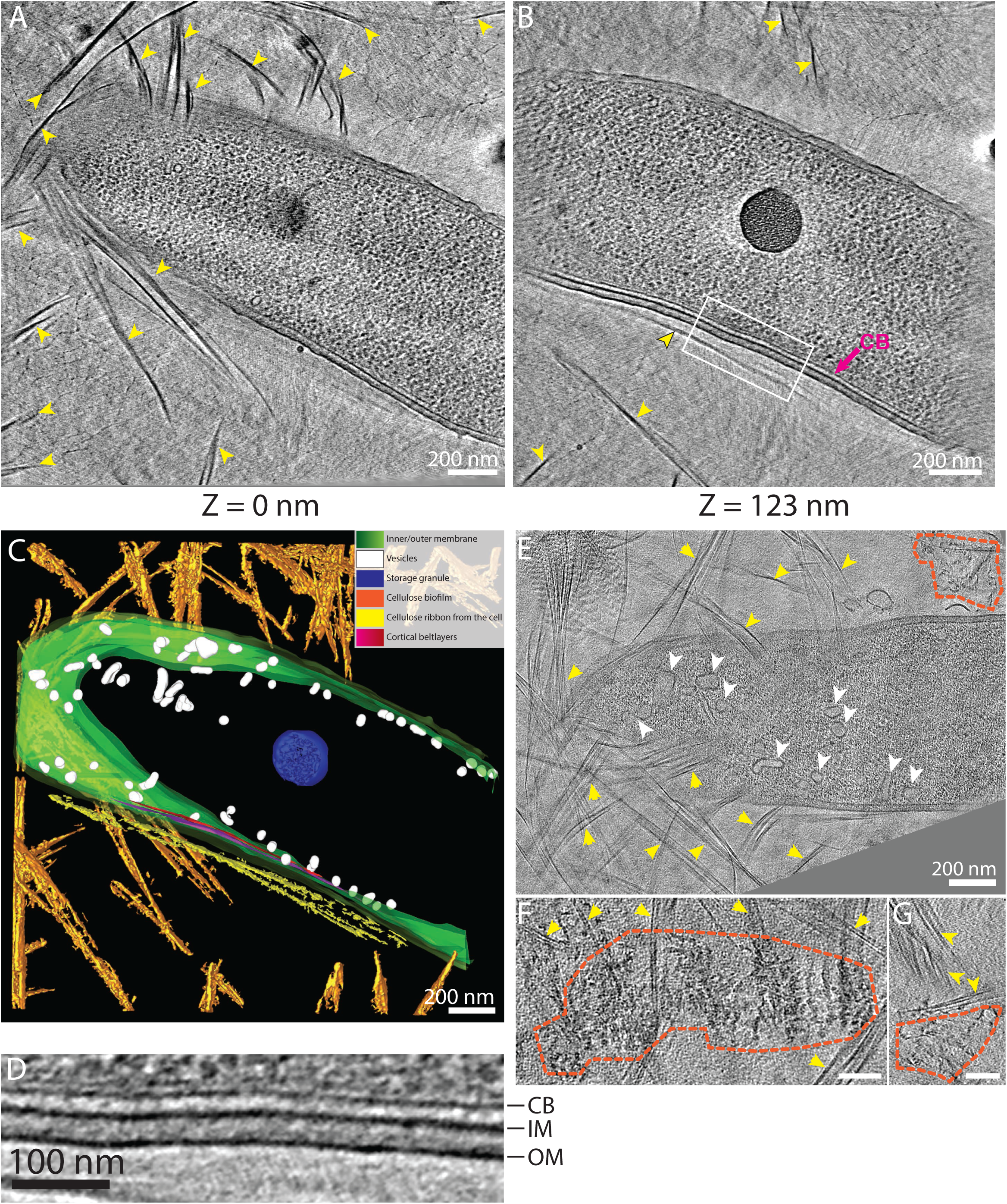
Lamellae of native biofilms also reveal numerous vesicles and the cortical belt. (**A-B**) Two tomographic slices of a *G. hansenii* cell from a biofilm grown for 6h surrounded by cellulose ribbons (yellow arrowheads). The cortical belt is visible in (B) (purple arrow) and seems to follow the trajectory of the cellulose sheet proximal to the OM (dark lined yellow arrowhead). (**C**) Manual segmentation of the tomogram displayed in (A) and (B) showing the juxtaposition of the cortical belt (purple to red) and the nascent cellulose ribbon (yellow). (**D**) Enlargement of the boxed region in (B) showing the layered cortical belt. (**E**) Tomographic slice of a cell surrounded by cellulose ribbons (yellow arrowheads) from a biofilm grown for 3h and harboring numerous vesicles in its cytosol (white arrowheads). Disorganized aggregates (orange dashed lines) are visible at this timepoint. (**F-G**) Tomographic slices showing additional examples of disorganized cellulose aggregates (orange dashed lines) surrounded by cellulose ribbons (yellow arrowheads) visible in 3h biofilms. Scale bars = 100 nm. All tomographic slices are 11-nm thick.

### The cortical belt is specific to bacterial species that produce crystalline cellulose ribbons

To see whether the cortical belt is specific to *G. hansenii*, we imaged another strain of *Gluconacetobacter, G. xylinus*, by cryo-ET at 300 minutes post-separation. Four out of 8 cells (50%) exhibited an extracellular cellulose ribbon along the cells’ long axis (Supplemental figure 3A). The cellulose ribbons observed had 2 sheets of cellulose, with an estimated average width of 27 ± 16 nm (n = 5). All four cells also possessed a cortical belt (Supplemental figure 3A-B, purple arrows), with similar dimensions to those in *G. hansenii*. The average distance from the cortical belt to the inner membrane was 24 ± 4 nm (n = 4). In one instance, the cortical belt also contained three stacked layers spaced (peak-to-peak) by 9 nm (Supplemental figure 3C). Aside from *Gluconacetobacter*, other bacterial species produce different types of cellulose. For instance, *Escherichia coli* 1094 can make amorphous cellulose^40^ and *Agrobacterium tumefaciens* makes crystalline cellulose microfibrils during plant infection^41^. Neither of these species are known to make cellulose ribbons. We asked whether structures similar to the cortical belt observed in *Gluconacetobacter* were present in these species. Our lab had previously imaged *A. tumefaciens* for other studies, and therefore cryo-tomograms of *A. tumefaciens* were already available. We confirmed by confocal microscopy that *A. tumefaciens* produces cellulose in the same growth conditions as had been used for the earlier experiments (Fig. 8A), and then screened the available tomograms for the presence of cellulose. As the purpose of the previous studies had not been cellulose synthesis observation, relatively few (65 out of 1,854 tomograms) showed distinct cellulose fibers in the vicinity of the cells (Fig. 8B-C, yellow arrowheads, supplemental video 4). These fibers did not adopt any preferential orientation and ran in all directions around the cell. They also had a decreased width (14 ± 5 nm, n = 52 fibers measured in 5 tomograms) compared to *G. hansenii* cellulose sheets, confirming that *A. tumefaciens* does not elaborate wide cellulose sheets nor ribbons but rather simpler structures of crystalline cellulose, presumably bundles of microfibrils. In the 65 cellulose-producing cells, we never observed a cortical belt structure. Two notable features were however observed: 1) a polar outer-membrane flattening in 28 cells with a thickening of the OM (43% out of the 65 cells presenting cellulose, Fig. 8B, cyan arrow) and 2) polar amorphous aggregates in 24 cells (37% out of the 65 cells presenting cellulose), (Fig. 8B, orange dashed lining). 19 cells exhibited all three described features, the polar flattening, the amorphous aggregates and the cellulose fibers. We suspect these polar amorphous aggregates to be the unipolar polysaccharides (UPP) described in previous work and shown to allow the attachment of *A. tumefaciens* to biotic and abiotic surfaces in the early stages of biofilm formation^42^. The very close proximity of the putative UPP to the polar flattening suggests the latter could hold the UPP-secreting complexes.

**Figure 8.**
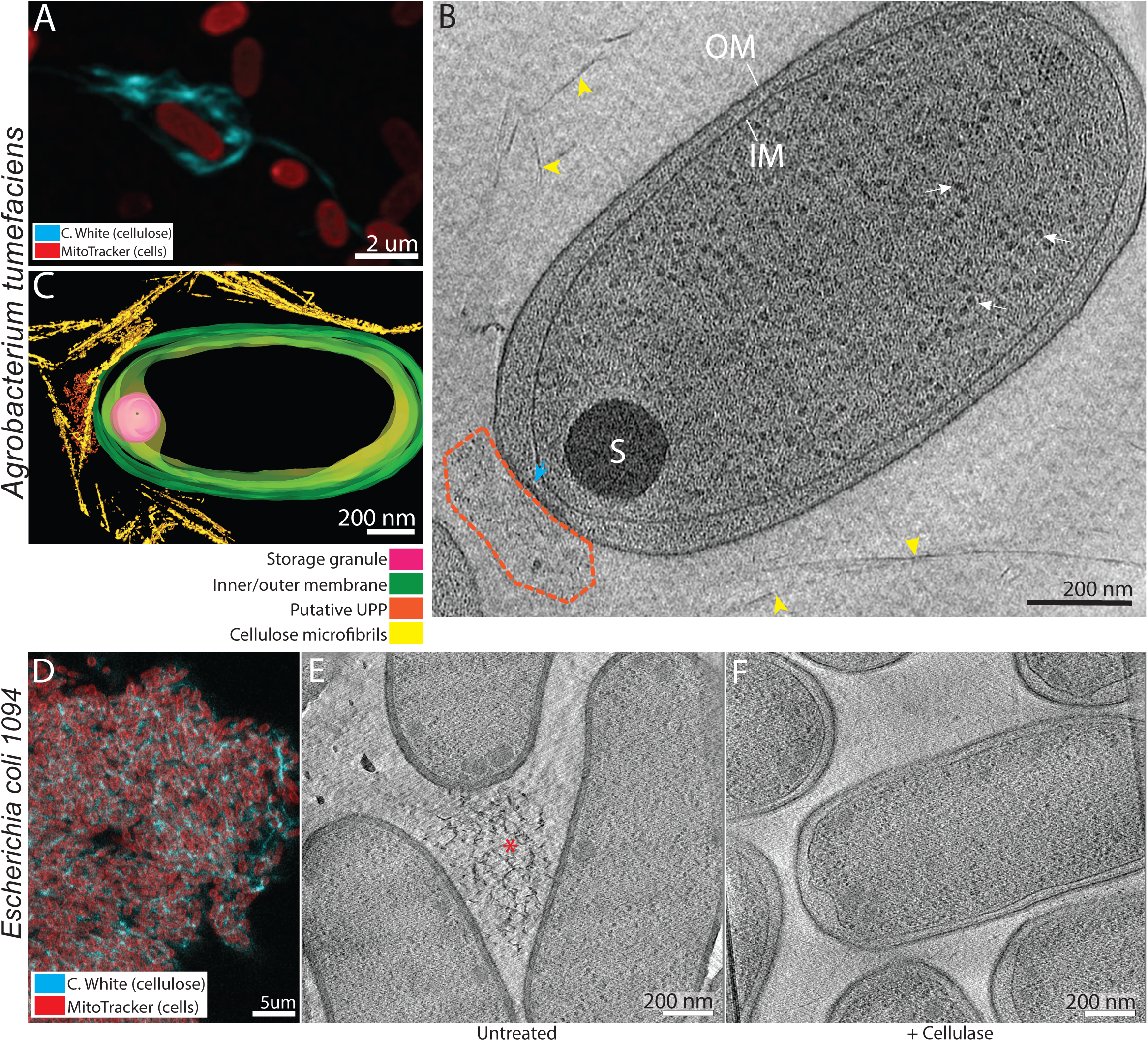
The cortical belt is not found in other cellulose-synthesizing species. (**A**) Maximum projection of *A. tumefaciens* cells synthesizing cellulose. Cells are stained with Mito Tracker Deep Red (red) and cellulose with Calcofluor-white (cyan). (**B**) 10-nm thick tomographic slice of a typical *A. tumefaciens* cell with cellulose microfibrils around (yellow arrowheads). No cortical belt can be seen in the cells. A polar flattening can be seen at the lower pole (cyan arrow) with an amorphous aggregate (orange dashed lines). These aggregates are most probably the UniPolar-Polysaccharide (UPP) synthesized specifically by *A. tumefaciens*. (**C**) Manual segmentation of the tomogram in (B) showing the organization of the cellulose microfibrils around the cell, the absence of the cortical belt and the putative UPP. (**D**) 50-nm optical slice of an induced *E. coli 1094* cellulose biofilm. Cells are stained with mitoTracker Deep Red (red) and cellulose with Calcofluor-white (cyan). (**E**) 6-nm tomographic slice of a lamella tomogram of bacterial mat showing three *E. coli 1094* cells and an amorphous cellulose aggregate between them (orange asterisk). (**F**) 6-nm tomographic slice of a lamella through a bacterial mat treated with cellulase, showing multiple cells. No cellulose was visible in this condition. No cortical belt can be seen in the cells in either condition.

We confirmed that *Escherichia coli* 1094 grown in minimal medium produces cellulose (Fig. 8D). The cells aggregated, making it difficult to image single cells by cryo-ET, so instead we FIB milled through bacterial mats, producing approximately 200 nm-thick lamellae. To identify cellulose structures, we also imaged lamellae from cultures grown in minimal medium supplemented with cellulase. In 3 of the 5 tomograms of untreated cells, we observed amorphous fibrous material (Fig. 8E, orange asterisk), that was not visible in 2 tomograms of a cellulase-treated culture (Fig. 8F). None of the cells imaged in either condition contained a cortical belt (n = 13 untreated and 5 cellulase-treated cells), suggesting that it is unique to bacteria producing higher-order crystalline cellulose structures, *i*.*e*. sheets.

## Discussion

Here we characterized bacterial cellulose synthesis in two *Gluconacetobacter* spp. and compared it to two other species by cryo-ET. We identified a novel cytoplasmic structure associated with the production of cellulose I ribbons in *Gluconacetobacter* spp. We also performed cryo-FIB milling followed by cryo-ET on a native biofilm.

### Cryo-ET confirms the need of a tight interaction between the nascent sheet and the OM

The cell-directed hierarchical model proposes linearly arranged 3.5-nm diameter pores on the surface of the cell^37^, each extruding an elementary fibril^23,29^. The arrangement of these pores in lines allows the crystallization of the elementary fibrils upon secretion and integration into a cellulose sheet parallel to the long axis of the cell^7,43,44^. Our results agree with this model. Indeed, we observed that when the gap between the nascent sheet and the OM exceeds approximately 40 nm, disorganized aggregates occur (Fig. 2). Along with previous work that observed similar events^23^, we hypothesize that these aggregates are microfibrils failing to integrate into an ordered sheet. Furthermore, it has been shown that the addition of compounds which bind directly to cellulose drastically alters the assembly of the sheets and leads to the formation of similar aggregates^23,44^. It appears as though preventing the nascent microfibrils from interacting with each other upon secretion prevents them from forming one organized sheet. Conversely, a confined spacing between the nascent sheet and the OM promotes proper crystallization of the nascent microfibrils. This proximity could be maintained either by a previously synthesized sheet preventing the nascent one from separating too far from the OM, or by specialized cellulose binding enzymes situated in the outer-leaflet of the OM, such as CmcAx, which has the ability to bind cellulose^45^.

### Cryo-ET sheds light on the buildup of a microfibril

Many values have been reported for the elementary fibrils’ dimension, mainly through direct observation by negative staining electron microscopy^14,29,36^. The most favored hypothesis is an approximately 1.5-nm thick elementary fibril (thoroughly discussed in ^23^). Very recently, the characterization of the structure of the BcsC subunit (the OM pore) strengthens this hypothesis, since the pore was seen to have an internal diameter of 1.5 nm at maximum^19^. Therefore, the simplest hypothesis is that each extrusion pore comprises a single BcsC sub-unit secreting an elementary fibril not more than 1.5 nm thick.

While negative staining has provided high-resolution views of cellulose ribbons ^23,36^, observing them in a frozen-hydrated state enables more accurate measurements of their dimensions and observation of their interaction with the OM. This is particularly important for extracellular polysaccharides, which have been shown to collapse and undergo drastic conformational changes upon dehydration, staining and embedding^46^.

We were able to image in two tomograms, microfibrils extruded perpendicularly to the OM and integrating to form a thin parallel sheet (Fig. 4E-F). A possible interpretation of why these events are rare is that they result from an accidental mechanical separation of the nascent sheet from the OM, revealing intermediate forms of cellulose bundling such as thin microfibrils. As explained earlier, precise measurement of the thickness of densities is difficult in cryo-ET since it is influenced by the defocus applied during imaging (causing overestimation of the true thickness). Despite this uncertainty, our measurements are done in a near native state. We estimated these microfibrils to be less than 11 nm in diameter (Fig. 4G-H). Previous work measured microfibril thicknesses from 3 to 12 nm in cellulose sheets splayed apart by cellulase treatments^47^. If we assume an elementary fibril is 1.5 nm in diameter since this is the maximal opening of a BcsC pore^19^, a 10 nm cylindrical microfibril would comprise approximately 35 elementary fibrils. This would require a cluster of 35 extrusion pores. Previous reports have stated the extrusion pores cluster in bunches from 2 to 4 pores^7,29^, substantially less than our calculation. Our data therefore suggest that there is more than one BcsC subunit per extrusion pore. For example, if each 3.5 nm diameter extrusion pore^37^ held 4 BcsC subunits, a cluster of 9 extrusion pores could produce a 9 nm diameter microfibril (Fig. 9). In this particular case, each extrusion pore holding multiple BcsC subunits would produce a crystalline aggregate of elementary fibrils which would pack with its neighboring aggregates to form a microfibril.

**Figure 9.**
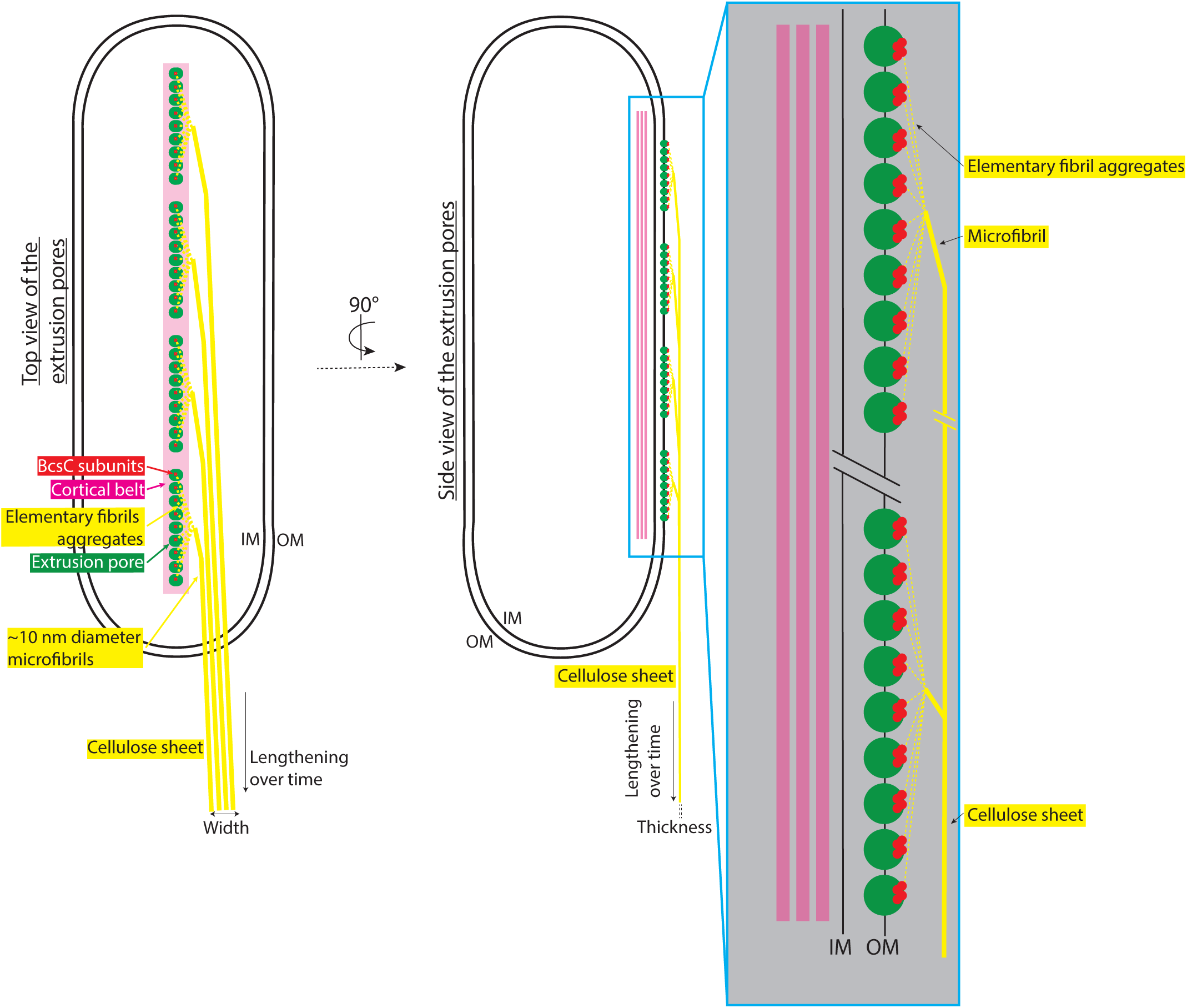
Updated cell-directed hierarchical model. Top (left) and side (right) view of a *G. hansenii* cell showing the different aggregation steps leading to a cellulose sheet, how microfibrils contribute to sheet width and the role of the cortical belt. In this model, clusters of 9 extrusion pores are depicted (green circles), the real numbers and distribution are unknown. Each extrusion pore is presented as comprising 4 BcsC subunits each (red circles), the actual number is not known. Inset in blue is a magnified view of the line of 9 extrusion pores, each hypothesized to extrude an aggregate of multiple elementary fibrils. All aggregates then coalesce to form a microfibril. These microfibrils then stack together, contributing to the width of the cellulose sheet. Adapted from the cell in figure 4E-G.

### Cryo-ET sheds light on the assembly of a cellulose Iα sheet

We found that ribbons were stacks of sheets that likely interact loosely with one another since the inter-sheet distance varied from 7- to 31-nm. This loose stacking corroborates previous observations^30^. Previous measurements done by negative staining had estimated cellulose sheet width to range from 40 to 600 nm^23,36,47^, wider than our measurements ranging from 11 to 69 nm (Fig. 4I). These variations have been attributed to the cell strain, growth conditions and intercellular variation^23,36,37^. We found that the thickness of cellulose sheets is similar to the diameter of the microfibrils. Therefore, our data suggest that microfibrils lie down in rows to create the width of the sheet. This was also suggested in^7^.

While the number of sheets produced by a single cell increased with time, the main dimension of growth appears to be ribbon length, as suggested by previous work and our fluorescence data showing cellulose ribbons several cell lengths long (Fig. 1A-B)^24^. Wider sheets occur in later time points (Fig. 4I), suggesting that sheet width also grows with time. However, in the current model, sheet width is correlated with the number of extrusion pores, hence to cell length^7,37^. It is possible that at 300min post-separation, cells are longer and possess more extrusion pores, therefore producing wider sheets.

### Cryo-ET on *G. hansenii* cells allowed the visualization of a novel cytoskeletal element, the cortical belt

Negative stain, cryo-fracture and immuno-EM studies have shown that cellulose extrusion pores in *Gluconacetobacter* align in a line on one side of the cell ^23,37,48^, but what causes this alignment is unknown. Here, we identify a novel cytoplasmic structure in two species of *Gluconacetobacter* that spatially correlates with the nascent cellulose ribbon (Fig. 1C-E and Fig. 5). This structure, which we term the cortical belt, is found at a fixed distance from the inner membrane (24 ± 4 nm) and remains intact upon cellulase treatment (Supplemental figure 1F, purple arrow), suggesting that it is stable even in the absence of the cellulose ribbon.

We observed the cortical belt in both *Gluconacetobacter* spp. imaged but not in other bacteria that produce less-ordered forms of cellulose, including *Escherichia coli* 1094, which synthesizes amorphous cellulose^40^, and *Agrobacterium tumefaciens*, which synthesizes cellulose I microfibrils^49^ (Fig. 8). This suggests that the cortical belt functions in the formation of cellulose I ribbons. BcsD and CcpAx, as well as two cell wall-related enzymes, have been shown to be involved in the crystallization process of the ribbons ^24,26,50^. It is possible that the cortical belt interacts with one or more of these components to guide the positioning of the BCS complexes, but the interaction may also be indirect. If the cortical belt is responsible for scaffolding the BCS complexes, it represents a novel prokaryotic cytoskeletal element, i.e. “a cytoplasmic protein filament and its associated superstructures that move or scaffold material within the cell”^51^. Other bacterial cytoskeletal elements have been observed to form belt-like structures, including bactofilins^52^, or to stack, like the CTP synthase^53^, although with different dimensions. *G. hansenii* has homologs of both bactofilin and CTP synthase, but neither shows co-evolution with the *bcs* gene cluster. We hope that future work will identify the component(s) that form the cortical belt, shedding more light on the molecular processes involved in the organization and clustering of the BCS complexes in *G. hansenii*.

### The cortical belt reveals another similarity between cellulose synthesis in *Gluconacetobacter* and land plants

Historically, the first plant cellulose synthase genes were identified by cDNA homology with the *G. xylinum* acsA (bcsA) gene^54^. Later on, phylogenetic studies highlighted an early divergence between cyanobacterial and plant cellulose synthases^55,56^. A large number of cellulose I synthesizing organisms have in common that the synthase complexes arrange in specific patterns, determining the final architecture of the cellulose structures^7^. A simple row in systems like *Gluconacetobacter* spp. or certain charophytes and chlorophytes^57^ and hexameric rosette structures called Cellulose Synthase Complexes (CSC) in land plants. In both, the extrusion of a crystalline form of cellulose exerts a force believed to be able to propel the CSCs in plants^58,59^ and the whole cell in *Gluconacetobacter*^24,36^. Our work uncovers an additional similarity, the involvement of a cytoskeletal element, the cortical belt, to guide the synthase complexes. In land plants CSCs have been shown to interact indirectly with underlying cortical microtubules, mediating trans-membrane cross-talk^60–62^, guiding and regulating CSC velocity^63–65^. While CSCs were shown to be motile in land plants, they are believed to be static in *Gluconacetobacter*^23^, perhaps held in place by the cortical belt, in order to transfer the propelling force to the whole cell.

### Insights from FIB-milling native biofilms

Cryo-FIB milling through native biofilms offers the possibility of observing bacteria in the context of their original biofilm environment and retrieving high resolution morphological and positional information about the cells relative to one another and relative to the biofilm layers. Visualization of the density and organization of the extracellular matrix and its interaction with the cells is also rendered possible by cryo-FIB milling. This is especially important since in nature most bacterial species are found in complex interacting communities, in the form of homogeneous or heterogeneous communities that organize in biofilms^10^.

Milling the *Gluconacetobacter* biofilms to 200 nm revealed numerous cytosolic vesicles of variable shapes and sizes. The cortical belt was also visible, as in the isolated cells. The cellulose ribbons aligned with each other to form larger arrays 2-3 µm wide (Fig. 6D, yellow arrowheads and supplemental video 3), showing the propensity of these structures to interact with each other. This propensity was previously characterized by live imaging of the cellulose biosynthesis and crystallization process in *Gluconacetobacter*, which showed that the bacterial cells preferentially follow already established tracks, *i.e*. previously synthesized cellulose ribbons^24^. The occurrence of disorganized cellulose clusters in biofilms grown for 3h but not 6h, suggests that such aggregates are either 1) digested by enzymes, likely CmcAx, reported to have an endoglucanase capable of digesting amorphous cellulose^66^ and to be present on the surface of *G. hansenii* or released in the environment^25,45^ or 2) diluted by a gradual increase in well-ordered ribbons over time.

Cell death in biofilms, with the fraction of dead cells measured at 10% in our biofilms, is a well-known phenomenon^10^, caused by programmed cell death mechanisms, cannibalistic behaviors such as already described in *B. subtilis*^67^ or nutrient/oxygen depletion^68,69^. We did not observe a preferential location of dead cells at the bottom of the biofilm, ruling out anoxic conditions being the primary cause of cell death. This could be because the thickness of the biofilm, between 1.5- and 3-um according to the cell depth distribution (Fig. 6H), is too small to have a significant oxygen gradient, as suggested by studies that measured total anoxia being reached generally between 70- and 80-um depth^69–71^. Processing thicker biofilms in the range of tens of microns would allow visualization of the effects of nutrient/oxygen gradients on cell distribution and physiology. For now, plunge freezing such as performed in this study can only properly vitrify samples less than ∼10 microns thick^72^. Moreover, milling thicknesses above 8-10 microns becomes labor intensive and technically difficult. A possible course of action for further studies would be to perform high-pressure freezing on thicker biofilms and then produce thin sections either by cryosectioning, hybrid cryosectioning/FIB-milling methods such as described in^73–75^ or following a cryo-lift out procedure^75^.

## Acknowledgments

This work was supported by NIH grant R35-GM122588 to GJJ, the Howard Hughes Medical Institute (HHMI) and the Center for Environmental Microbial Interactions (CEMI) pilot grant program. Cryo-electron microscopy was performed in the Beckman Institute Resource Center for Transmission Electron Microscopy at Caltech. We thank Jean Marc Ghigo for kindly providing us the E. Coli 1094 strain. Special acknowledgments to Catherine Oikonomou for all the help and scientific advice given during this study and also to Candace Haigler for sharing her thoughts and her precious experience on the not so common *Gluconacetobacter* spp.

## Methods

### Cell culture

*Gluconacetobacter hansenii* (ATCC 23769) was cultured as previously described ^35^ in SH medium: 2% glucose, 0.5% bactopeptone, 0.5% yeast extract, pH 6. For solid medium, 2.5% bacto-agar was added. Cells were separated from the cellulose biofilm by mechanical disruption as previously described^36^. Briefly, the bacterial cellulose biofilm developing at the air-media interface was picked up with a single-use sterile inoculating loop and transferred to fresh medium, where it was vigorously shaken and then removed. In preparation for freezing, cells were pelleted by centrifugation for 10 minutes at 2500rcf at 20C and resuspended in 0.5mL of SH media. The culture was incubated for the desired length of time at 30°C without shaking before plunge freezing. For cellulose digestion, 0.2g/L cellulase (Worthington, purified exo- and endo-glucanases, #LS002598) was added.

*Gluconacetobacter xylinus* (ATCC 700178/BPR2001) was cultured as described above in Fructose–Peptone–Yeast Extract (FPY) media: 2% fructose, 1% bactopeptone, 0.5% yeast extract and 0.25% K_2_HPO_4_.

*Escherichia coli* 1094 was cultured in Lysogeny Broth (LB) and induced for cellulose production in minimal medium: 0.2% (NH_4_)_2_SO_4_, 1.4% KH_2_PO_4_, 0.1% MgSO_4_, 0.5% FeSO_4_.7H_2_O, 0.4% glucose, 0.01% thiamine, pH 7. A saturated overnight LB culture was diluted 1:50 into 3mL of minimal medium with or without 0.2g/L cellulase (Worthington, purified exo- and endo-glucanases, #LS002598). Cultures were incubated at 37°C with shaking at 220rpm. When the medium transitioned from turbid to clear and white flakes appeared (cellulose and bacteria), the induction of cellulose synthesis is considered successful.

*Agrobacterium tumefaciens* was cultured as described in previous work^76^. Briefly, *A. tumefaciens* C58 was cultivated in liquid AB medium (glucose 0.2%, NH4Cl 18.7mM, MgSO4 2.5uM, KCl 2mM, CaCl2 0.07mM, FeSO4 0.01mM, K2HPO4 8.4mM, NaH2PO4.7H2O 4.16mM, pH 7) at 30C overnight. Induction was done by pipetting 100uL of overnight culture and spreading onto AB induction plates (glucose 0.2%, NH4Cl 18.7mM, MgSO4 2.5uM, KCl 2mM, CaCl2 0.07mM, FeSO4 0.01mM, K2HPO4 8.4mM, NaH2PO4.7H2O 4.16mM, Bactagar 1.7%, Acetosyringone 100uM, pH 5.8). Plates were then incubated for 3 days at 20C. Cells were resolubilized by scraping a small amount from the plate with an inoculation loop and resuspending it in 100uL of liquid induction AB medium.

The following strains are the ones included in the tomogram analysis: NT1 is a C58 strain without the pTiC58 (tumor inducing) plasmid; A139 strain is NT1REB(pJK270) + pJZ041. NT1REB is a “bald strain”, no flagellin mutant, derived from NT1. The pJK270 is pTiC58 with the transposed NPTII gene for kanamycin resistance. The pJZ041 plasmid carries a GFP tagged VirB8 gene, a component of the T4SS (Aguilar et al. 2011); JX148 strain is a C58 derived mutant of the rem gene. The strain is non motile; AD348 is a GV3101(pMP90) strain with its whole VirB system deleted. GV3101 is a pTiC58 free, rifampicin resistant C58 strain and pMP90 is a helper pTiC58 without the T-DNA; AD1484 is a AD348 variant, transformed with pAD2079 containing the whole VirB system.

### Confocal microscopy

Cellulose was stained with Calcofluor-white (Sigma-Aldrich, #18909) at a concentration of 0.001% and cell membranes were stained with MitoTracker Deep Red FM (Thermo-Fisher, #M22426) at a concentration of 0.5ug/uL. Stack acquisition was done on a Zeiss LSM880 Airy Scan microscope. Airy scan acquisitions were performed in super-resolution mode with Z-step set at the optimal optical sectioning. The Mito-Tracker Deep Red FM channel was set as the following: excitation at 633 nm, use of the 488/561/633 main beam splitter and a band-pass 570-620 + long-pass 645 filter. The Calcofluor White channel was set as the following: excitation at 405 nm, use of the 405 main beam splitter and a band-pass 420-480 + band-pass 495-550 filter. Airy scan processing was performed on the fly by the in-built algorithm of Zeiss Black.

### Sample preparation for cryo-EM

For isolated cells, Quantifoil Cu R2/2 Finder grids (*Quantifoil Micro Tools GmbH*) were glow-discharged at 15mA for 1min. The grids were pre-incubated with fiducial marker solution prepared as follows: 50µL of 10nm colloidal gold (*Ted Pella, Inc*) mixed with 50uL of 5% BSA, vortexed 1 min and centrifuged at 15,000rcf for 15 min, supernatant discarded, and pellet resuspended in 40µL of PBS buffer. 3μL were deposited on each grid, left for 1 minute then back-blotted with Whatman paper. Cells were plunge frozen with a Vitrobot Mark IV (*Thermo Fisher Scientific*) with 100% humidity at 30°C and back-blotted for 3 to 5s.

For native biofilms, Quantifoil gold R2/2 Finder grids were placed in 35mm glass bottom petri dishes (*MatTek Corporation* #P35G-1.0-2.0C) containing 1mL of SH media inoculated with a 2-day old biofilm. The dishes were sealed with Micropore tape (*3M*) and incubated without shaking at 30°C for 3 to 6 hours. Plunge-freezing was done at 22C, 50% humidity, either with manual blotting on both sides of the grids (first back-blotted then front-blotted) or using the automatic blotting function of the Vitrobot with a blot time of 5-6s, blot force of 15 and drain time of 2s.

For *E. coli* 1094, after 4 hours of incubation in minimal media, the medium should turn from turbid to clear with white flakes. OD_600_ of the cultures was monitored using the culture (always turbid) where cellulose induction was performed in the presence of cellulase to keep the cells from aggregating. It was then used as a reference to concentrate the cells to high OD_600_ (10-20), in order to form bacterial mats on the EM grids, for control and cellulase conditions. Plunge-frozen was done at 20C, 100%, either with manual back-blotting for 5-7s and a drain time of 1s or using the automatic blotting function of the Vitrobot with a wait time of 10s, blot time of 5-6s, blot force of 3 and drain time of 1s.

### FIB milling

Grids were clipped in Autogrid holders (*Thermo Fisher*) machined with a notch to allow FIB milling closer to the edge of the grid. Autogrids were placed in a custom-built shuttle and inserted into a Versa 3D dual-beam FIB/SEM microscope with FEG (*FEI*) equipped with a PP3000T cryo-transfer apparatus (*Quorum Technologies*). They were maintained at −175°C at all times by a custom-built cryo-stage^77^. To reduce sample charging and protect the sample from curtaining during milling, the grids were sputter-coated with platinum at 15 mA for 60 seconds. Thin lamellae were generated with the Ga^+^ ion beam at 30 kV at angles ranging from 10 to 17 degrees. Rough milling was done at high currents, ranging from 0.3 nA to 100 pA until the lamellae measured 1 micron in thickness under the FIB view. Current was then progressively brought down to 10 pA for the final milling steps until the measured thickness was between 100-200 nm. Final polishing of the back end of the lamella is also done at 10pA where the sample is tilted +0.5 to 1° to homogenize the lamella thickness. During the whole procedure, imaging with the SEM beam was done at 5 kV and 13 pA.

### Electron cryo-tomography

Tomography of whole cells and FIB-milled lamellae was performed on either a Titan Krios or Tecnai G2 Polara transmission electron microscope (*Thermo Fisher*) equipped with 300 kV field emission gun, energy filter (*Gatan*) and K2 or K3 Summit direct electron detector (*Gatan*). The Krios is equipped with a Volta phase plate (*Thermo Fisher*) ^78^. Tilt-series acquisition was done with SerialEM ^79^ with a 2-3° tilt increment for a total range of ±60° or ±50°, defocus of −4, −6 or −8 µm, and total dose up to 180 e^-^/Å^2^. Volta phase plate imaging was performed in Figures 1, 2, 5 and 7A-B with a defocus of −2µm and a measured phase shift of 0.5 π/rad before tilt series acquisitions.

Low magnification tomography on the biofilm lamellae was performed at 6500 magnification (14 Å^2^ pixel size) with a −10 or −15 μm defocus and a total dose between 5 and 10 e^-^/Å^2^. Tomography of FIB-milled lamellae was done exclusively on the Titan Krios. Because samples were thinner, the total dose was limited to ∼80 e^-^/Å^2^.

### Data processing

Tomograms were reconstructed using the IMOD software (http://bio3d.colorado.edu/imod/) ^80^. Alignment was done on 1k x 1k binned tilt-series with fiducial-based alignment. Aligned stacks were low-pass filtered (0.35, σ = 0.05) to eliminate high-frequency noise. Weighted back projection reconstruction was performed and the “SIRT-like filter” was used with 20 iterations. Segmentation was also done using IMOD and drawing tools developed by Andrew Noske (http://www.andrewnoske.com/student/imod.php). To better distinguish features during the segmentation steps, tomograms were filtered with the 3D non-linear anisotropic diffusion filter in IMOD. The cell contours and cortical belt were segmented manually on a Cintiq 21uX tablet (*Wacom*) and cellulose was segmented using a semi-automated thresholded method. 1) A denoising Non-linear Anisotropic Diffusion filter was applied (included in the *etomo* package, http://bio3d.colorado.edu/imod/) on the tomogram; 2) Precise boundary models are drawn around the structures to be thresholded; 3) Thresholding segmentation is performed with 3Dmod using the isosurface function and the previously drawn contours are used as a mask. When the contours are precisely following the contours, this technic allows to raise the isosurface threshold without picking up background noise.

Measurements for all distances between elements (cellulose sheet – outer-membrane, width of the cellulose ribbon, cortical belt – inner-membrane) were taken by generating normalized density profile plots and measuring the distances between the density peaks of the corresponding sub-cellular features (Fig. 3). This was automated with a custom script, *<u>sideview-profile-average</u>*, written by Davi Ortega (https://www.npmjs.com/package/sideview-profile-average).

Estimation of the cell depth in the native biofilm lamellae was calculated as follows: 1) using the two parallel walls of the milled trench, a perpendicular line is traced at the leading edge of the lamella (where the platinum meets the frozen material); 2) Lines are drawn from the center of the cells to the leading edge perpendicular line (Fig. 6H, red line in top view of lamella); 3) The distance from the cell center to the limit of the platinum on the leading edge, which is the surface of the sample, is measured. The real depth is then calculated using the following equation: opposite side (real depth) = tan (*a*) x adjacent side (distance measured, d in Fig. 6H). The angle *a* is the angle between the grid surface and the FIB gun during the milling process, which can be accurately measured during reconstruction with 3dmod.

### Statistical analysis

All statistics were performed with GraphPad Prism software (https://www.graphpad.com/scientific-software/prism/). All datasets were first analyzed for normality using the Shapiro-Wilk test and homoscedasticity (equal standard deviations). If dataset is normal, appropriate parametric tests were performed and if not, appropriate non-parametric tests were performed.

**Figure 2E:** n = 3 and 24 for the “loose” and “tight” configuration respectively. Two tailed P-value = 0.0008, Mann-Whitney test.

**Figure 4A**: n = 6, 15, 33 for 13-, 20- and 300-minutes, respectively.

**Figure 4B**: n = 6 and n = 21 tomograms for 20- and 300-minutes post-separation, respectively. Two tailed P-value < 0.0001, One sample Wilcoxon signed rank test against a theoretical value of 1 (number of sheets observed at 20-min post-separation).

**Figure 4H**: n = 12 and 4 microfibril thickness measurements performed on two separate tomograms (Cell #1 and #2, left side of the graph). N = 47 measurements for inter-sheet distances performed on 23 tomograms. ANOVA followed by Tukey’s multiple comparison test was performed. Cell #1 vs Cell #2, Cell #1 vs 300-min inter-sheet distances and Cell #2 vs 300-min inter-sheet distances showed adjusted P-values of 0.073, 0.15 and 0.0015, respectively.

**Figure 4I**: n = 6 and 45 sheets measured at 20- and 300-minutes post-separation. Welch’s t test (parametric t-test without equal SD assumption) showed a P-value of 0.23.

**Figure 6F**: n = 6 and 4 for biofilms let to grow for 3h and 6h, respectively. Unpaired T-test showed a two-tailed P-value of 0.0011.

**Figure 6G**: n = 6 and 4 for biofilms let to grow for 3h and 6h, respectively. Unpaired T-test showed a two-tailed P-value of 0.2720.

**Figure 6H**: n = 49, 46, 4 and 11 for live and dead cells in 3h and 6h biofilms, respectively. Mann-Whitney tests were performed on live vs dead cells in 3h and 6h biofilms conditions, showing two-tailed P-values of 0.82 and 0.54, respectively.

## Supplemental movie legends

**Supplemental video 1** | Movie showing multiple views of the tomographic volume shown in Figure 1C-E. The disposition of the cellulose ribbon relative to the cell and its very close contact with the bacterial envelope is demonstrated.

**Supplemental video 2** | Movie showing multiple views of the tomographic volume shown in Figure 5A-C. The close association between the cortical belt and the cellulose ribbon are shown. The second part of the animation shows the tomographic volume shown in Figure 2 and Figure 5D-F. The multilayered structure of the cortical belt is visible.

**Supplemental video 3** | Movie of the FIB-milling of a lamella through the *G. hansenii* biofilms shown in Figure 6 and the 3D organization of the cells and the cellulose within the biofilm shown in Figure 6. The second part of the animation shows the tomographic volume shown in Figure 7A-D. Numerous cytoplasmic vesicles and the cortical belt underneath a cellulose ribbon are visible.

**Supplemental video 4** | Movie showing multiple views of the tomographic volume shown in Figure 8B-C. The cellulose fibers are seen around the cell. No cortical belt is visible. A polar flattening is visible at the top of the cell with a “thickening” of the OM and a large periplasmic density underneath. The putative UPP (amorphous aggregate) is seen on the side of the polar flattening.

## Bibliography

1. Pauly, Markus & Keegstra, Kenneth. Cell-wall carbohydrates and their modification as a resource for biofuels. Plant Journal vol. 54 559–568 (2008).

2. Pauly, Markus & Keegstra, Kenneth. Plant cell wall polymers as precursors for biofuels. Current Opinion in Plant Biology vol. 13 305–312 (2010).

3. Hon, David N. S. Cellulose: a random walk along its historical path. Cellulose 1, 1–25 (1994).

4. Xu, Youjie, Zhang, Meng, Roozeboom, Kraig & Wang, Donghai. Integrated bioethanol production to boost low-concentrated cellulosic ethanol without sacrificing ethanol yield. Bioresour. Technol. 250, 299–305 (2018).

5. Gatenholm, Paul & Klemm, Dieter. Bacterial Nanocellulose as a Renewable Material for Biomedical Applications. MRS Bull. 35, 208–213 (2010).

6. Cavalcante, Aline Ribeiro Teixeira, Lima, Rodrigo Pontes de, Souza, Veridiana Sales Barbosa de, Pinto, Flávia Cristina Morone, Campos Júnior, Olavio, Silva, Jaiurte Gomes Martins da, Albuquerque, Amanda Vasconcelos de & Aguiar, José Lamartine de Andrade. Effects of bacterial cellulose gel on the anorectal resting pressures in rats submitted to anal sphincter injury. Heliyon 4, e01058 (2018).

7. Brown Jr., R. M. The biosynthesis of cellulose. J. Macromol. Sci. Part A Pure Appl. Chem. 33, 1345–1373 (1996).

8. McNamara, Joshua T., Morgan, Jacob L. W. & Zimmer, Jochen. A molecular description of cellulose biosynthesis. Annu. Rev. Biochem. 84, 895–921 (2015).

9. Cosgrove, Daniel J. Growth of the plant cell wall. Nat. Rev. Mol. Cell Biol. 6, 850–861 (2005).

10. Flemming, Hans Curt, Wingender, Jost, Szewzyk, Ulrich, Steinberg, Peter, Rice, Scott A. & Kjelleberg, Staffan. Biofilms: An emergent form of bacterial life. Nature Reviews Microbiology vol. 14 563–575 (2016).

11. Römling, Ute & Galperin, Michael Y. Bacterial cellulose biosynthesis: Diversity of operons, subunits, products, and functions. Trends in Microbiology vol. 23 545–557 (2015).

12. De Vos, Willem M. Microbial biofilms and the human intestinal microbiome. npj Biofilms and Microbiomes vol. 1 15005 (2015).

13. Costerton, J. W., Cheng, K. J., Geesey, G. G., Ladd, T. I., Nickel, J. C., Dasgupta, M. & Marrie, T. J. Bacterial Biofilms in Nature and Disease. Annu. Rev. Microbiol. 41, 435–464 (1987).

14. Haigler, Candace Hope, Brown, R. Malcolm & Benziman, Moshe. The fine structure of cellulose microfibrils. Science 119, 80–82 (1980).

15. Morgan, Jacob L. W., McNamara, Joshua T., Fischer, Michael, Rich, Jamie, Chen, Hong Ming, Withers, Stephen G. & Zimmer, Jochen. Observing cellulose biosynthesis and membrane translocation in crystallo. Nature 531, 329–334 (2016).

16. Du, Juan, Vepachedu, Venkata, Cho, Sung Hyun, Kumar, Manish & Nixon, B. Tracy. Structure of the cellulose synthase complex of Gluconacetobacter hansenii at 23.4 Å resolution. PLoS One 11, e0155886 (2016).

17. Hu, S. Q., Gao, Y. G., Tajima, Kenji, Sunagawa, Naoki, Zhou, Yong, Kawano, Shin, Fujiwara, Takaaki, Yoda, Takanori, Shimura, Daisuke, Satoh, Yasuharu, Munekata, Masanobu, Tanaka, Isao & Yao, Min. Structure of bacterial cellulose synthase subunit D octamer with four inner passageways. Proc. Natl. Acad. Sci. 107, 17957–17961 (2010).

18. Saxena, I. M., Kudlicka, K., Okuda, K. & Brown, R. M. Characterization of genes in the cellulose-synthesizing operon (acs operon) of Acetobacter xylinum: Implications for cellulose crystallization. J. Bacteriol. 176, 5735–5752 (1994).

19. Acheson, Justin F., Derewenda, Zygmunt S. & Zimmer, Jochen. Architecture of the Cellulose Synthase Outer Membrane Channel and Its Association with the Periplasmic TPR Domain. Structure (2019).

20. Whitney, John C., Hay, Iain D., Li, Canhui, Eckford, Paul D. W., Robinson, Howard, Amaya, Maria F., Wood, Lynn F., Ohman, Dennis E., Bear, Christine E., Rehm, Bernd H. & Lynne Howell, P. Structural basis for alginate secretion across the bacterial outer membrane. Proc. Natl. Acad. Sci. 108, 13083–13088 (2011).

21. Rehman, Zahid U., Wang, Yajie, Moradali, M. Fata, Hay, Iain D. & Rehm, Bernd H. A. Insights into the assembly of the alginate biosynthesis machinery in Pseudomonas aeruginosa. Appl. Environ. Microbiol. 79, 3264–3272 (2013).

22. Keiski, Carrie-Lynn, Harwich, Michael, Jain, Sumita, Neculai, Ana Mirela, Yip, Patrick, Robinson, Howard, Whitney, John C., Riley, Laura, Burrows, Lori L., Ohman, Dennis E. & Howell, P. Lynne. AlgK Is a TPR-Containing Protein and the Periplasmic Component of a Novel Exopolysaccharide Secretin. Structure 18, 265–273 (2010).

23. Haigler, Candace Hope. Alteration of cellulose assembly in Acetobacter xylinum by fluorescenet brightening agents, direct dyes and cellulose derivatives. (University of North Carolina, 1982).

24. Mehta, Kalpa, Pfeffer, Sarah & Brown, R. Malcolm. Characterization of an acsD disruption mutant provides additional evidence for the hierarchical cell-directed self-assembly of cellulose in Gluconacetobacter xylinus. Cellulose 22, 119–137 (2015).

25. Nakai, Tomonori, Sugano, Yasushi, Shoda, Makoto, Sakakibara, Hitoshi, Oiwa, Kazuhiro, Tuzi, Satoru, Imai, Tomoya, Sugiyama, Junji, Takeuchi, Miyuki, Yamauchi, Daisuke & Mineyukia, Yoshinobu. Formation of highly twisted ribbons in a carboxymethylcellulase gene-disrupted strain of a cellulose-producing bacterium. J. Bacteriol. 195, 958–964 (2013).

26. Sunagawa, Naoki, Fujiwara, Takaaki, Yoda, Takanori, Kawano, Shin, Satoh, Yasuharu, Yao, Min, Tajima, Kenji & Dairi, Tohru. Cellulose complementing factor (Ccp) is a new member of the cellulose synthase complex (terminal complex) in Acetobacter xylinum. J. Biosci. Bioeng. 115, 607–612 (2013).

27. Deng, Ying, Nagachar, Nivedita, Xiao, Chaowen, Tien, Ming & Kao, Teh Hui. Identification and characterization of non-cellulose-producing mutants of Gluconacetobacter hansenii generated by Tn5 transposon mutagenesis. J. Bacteriol. 195, 5072–5083 (2013).

28. Cousins, Susan K. & Brown, R. Malcolm. Cellulose I microfibril assembly: computational molecular mechanics energy analysis favours bonding by van der Waals forces as the initial step in crystallization. Polymer 36, 3885–3888 (1995).

29. Haigler, Candace H. & Benziman, Moshe. Biogenesis of Cellulose I Microfibrils Occurs by Cell-Directed Self-Assembly in Acetobacter xylinum. in Cellulose and Other Natural Polymer Systems 273–297 (Springer US, 1982).

30. Cousins, Susan K. & Brown, R. Malcolm. Photoisomerization of a dye-altered β-1,4 glucan sheet induces the crystallization of a cellulose-composite. Polymer 38, 903–912 (1997).

31. Saxena, I. M. & Brown, R. M. Identification of a second cellulose synthase gene (acsAII) in Acetobacter xylinum. J. Bacteriol. 177, 5276–83 (1995).

32. Florea, Michael, Reeve, Benjamin, Abbott, James, Freemont, Paul S. & Ellis, Tom. Genome sequence and plasmid transformation of the model high-yield bacterial cellulose producer Gluconacetobacter hansenii ATCC 53582. Sci. Rep. 6, 23635 (2016).

33. Toyosaki, Hiroshi, Kojima, Yukiko, Tsuchida, Takayasu, Hoshino, Ken-Ichiro, Yamada, Yuzo & Yoshinaga, Fumihiro. The characterization of an acetic acid bacterium useful for producingbacterial cellulose in agitation cultures: The proposal of Acetobacter xylinum subsp. sucrofermentans subsp. nov. J. Gen. Appl. Microbiol. 41, 307–314 (1995).

34. Park, Joong Kon, Jung, Jae Yong & Park, Youn Hee. Cellulose production by Gluconacetobacter hansenii in a medium containing ethanol. Biotechnol. Lett. 25, 2055–2059 (2003).

35. Schramm, M. & Hestrin, S. Factors affecting Production of Cellulose at the Air/ Liquid Interface of a Culture of Acetobacter xylinum. J. Gen. Microbiol. 11, 123–129 (1954).

36. Brown, R. M., Willison, J. H., Richardson, C. L. & Richardson, C. L. Cellulose biosynthesis in Acetobacter xylinum: visualization of the site of synthesis and direct measurement of the in vivo process. Proc. Natl. Acad. Sci. U. S. A. 73, 4565–9 (1976).

37. Zaar, K. Visualization of pores (export sites) correlated with cellulose production in the envelope of the gram-negative bacterium Acetobacter xylinum. J. Cell Biol. 80, 773–777 (1979).

38. Hawkes, Peter W. The electron microscope as a structure projector. in Electron Tomography: Methods for Three-Dimensional Visualization of Structures in the Cell vol. 9780387690087 83–111 (Springer New York, 2006).

39. Radermacher, Michael. Weighted back-projection methods. in Electron Tomography: Methods for Three-Dimensional Visualization of Structures in the Cell vol. 9780387690087 245–273 (Springer New York, 2006).

40. Le Quéré, Benjamin & Ghigo, Jean Marc. BcsQ is an essential component of the Escherichia coli cellulose biosynthesis apparatus that localizes at the bacterial cell pole. Mol. Microbiol. 72, 724–740 (2009).

41. Matthysse, A. G., Holmes, K. V & Gurlitz, R. H. G. Elaboration of cellulose fibrils by Agrobacterium tumefaciens during attachment to carrot cells. J. Bacteriol. 145, 583–595 (1981).

42. Xu, Jing, Kim, Jinwoo, Koestler, Benjamin J., Choi, Jeong Hyeon, Waters, Christopher M. & Fuqua, Clay. Genetic analysis of agrobacterium tumefaciens unipolar polysaccharide production reveals complex integrated control of the motile-to-sessile switch. Mol. Microbiol. 89, 929–948 (2013).

43. Ross, Peter, Mayer, Raphael, Benziman, A. N. D. Moshe & Benziman, M. Cellulose Biosynthesis and Function in Bacteria. Microbiology 55, 35–58 (1991).

44. Benziman, Moshe, Haigler, Candace H., Brown, R. Malcolm, White, Alan R. & Cooper, Kay M. Cellulose biogenesis: Polymerization and crystallization are coupled processes in Acetobacter xylinum. Proc. Natl. Acad. Sci. U. S. A. 77, 6678–6682 (1980).

45. Yasutake, Yoshiaki, Kawano, Shin, Tajima, Kenji, Yao, Min, Satoh, Yasuharu, Munekata, Masanobu & Tanaka, Isao. Structural characterization of the Acetobacter xylinum endo-β-1,4-glucanase CMCax required for cellulose biosynthesis. Proteins Struct. Funct. Bioinforma. 64, 1069–1077 (2006).

46. Dohnalkova, Alice C., Marshall, Matthew J., Arey, Bruce W., Williams, Kenneth H., Buck, Edgar C. & Fredrickson, James K. Imaging hydrated microbial extracellular polymers: Comparative analysis by electron microscopy. Appl. Environ. Microbiol. 77, 1254–1262 (2011).

47. White, Alan R. & Brown, R. M. Enzymatic hydrolysis of cellulose: Visual characterization of the process. Proc. Natl. Acad. Sci. U. S. A. 78, 1047–1051 (1981).

48. Kimura, S., Chen, H. P., Saxena, I. M., Brown, J. & Itoh, T. Localization of c-di-GMP-binding protein with the linear terminal complexes of Acetobacter xylinum. J. Bacteriol. 183, 5668–5674 (2001).

49. Matthysse, Ann G., White, Sally & Lightfoot, Richard. Genes required for cellulose synthesis in Agrobacterium tumefaciens. J. Bacteriol. 177, 1069–1075 (1995).

50. Deng, Ying, Nagachar, Nivedita, Fang, Lin, Luan, Xin, Catchmark, Jeffrey M., Tien, Ming & Kao, Teh Hui. Isolation and characterization of two cellulose morphology mutants of Gluconacetobacter hansenii ATCC23769 producing cellulose with lower crystallinity. PLoS One 10, e0119504 (2015).

51. Pilhofer, Martin & Jensen, Grant J. The bacterial cytoskeleton: More than twisted filaments. Current Opinion in Cell Biology vol. 25 1–9 (2013).

52. Kühn, Juliane, Briegel, Ariane, Mörschel, Erhard, Kahnt, Jörg, Leser, Katja, Wick, Stephanie, Jensen, Grant J. & Thanbichler, Martin. Bactofilins, a ubiquitous class of cytoskeletal proteins mediating polar localization of a cell wall synthase in Caulobacter crescentus. EMBO J. 29, 327–339 (2010).

53. Ingerson-Mahar, Michael, Briegel, Ariane, Werner, John N., Jensen, Grant J. & Gitai, Zemer. The metabolic enzyme CTP synthase forms cytoskeletal filaments. Nat. Cell Biol. 12, 739–746 (2010).

54. Pear, J. R., Kawagoe, Y., Schreckengost, W. E., Delmer, D. P. & Stalker, D. M. Higher plants contain homologs of the bacterial celA genes encoding the catalytic subunit of cellulose synthase. Proc. Natl. Acad. Sci. U. S. A. 93, 12637–42 (1996).

55. Nobles, David R. & Brown, R. Malcolm. The pivotal role of cyanobacteria in the evolution of cellulose synthases and cellulose synthase-like proteins. Cellulose 11, 437–448 (2004).

56. Nobles, D. R., Romanovicz, D. K. & Brown, Jr. Cellulose in cyanobacteria. Origin of vascular plant cellulose synthase? Plant Physiol. 127, 529–542 (2001).

57. Lampugnani, Edwin R., Flores-Sandoval, Eduardo, Tan, Qiao Wen, Mutwil, Marek, Bowman, John L. & Persson, Staffan. Cellulose Synthesis – Central Components and Their Evolutionary Relationships. Trends in Plant Science vol. 24 402–412 (2019).

58. Diotallevi, Fabiana & Mulder, Bela. The cellulose synthase complex: A polymerization driven supramolecular motor. Biophys. J. 92, 2666–2673 (2007).

59. Chan, Jordi, Coen, Enrico, Chan, Jordi & Coen, Enrico. Interaction between Autonomous and Microtubule Guidance Systems Controls Cellulose Synthase Report Interaction between Autonomous and Microtubule Guidance Systems Controls Cellulose Synthase Trajectories. Curr. Biol. 1–7 (2020).

60. Paredez, Alexander R., Somerville, Christopher R. & Ehrhardt, David W. Visualization of cellulose synthase demonstrates functional association with microtubules. Science 312, 1491–1495 (2006).

61. Li, Shundai, Lei, Lei, Somerville, Christopher R. & Gu, Ying. Cellulose synthase interactive protein 1 (CSI1) mediates the intimate relationship between cellulose microfibrils and cortical microtubules. Plant Signal. Behav. 7, 1–5 (2012).

62. Sampathkumar, Arun, Peaucelle, Alexis, Fujita, Miki, Schuster, Christoph, Persson, Staffan, Wasteneys, Geoffrey O. & Meyerowitz, Elliot M. Primary wall cellulose synthase regulates shoot apical meristem mechanics and growth. Development 146, (2019).

63. Fujita, Miki, Himmelspach, Regina, Ward, Juliet, Whittington, Angela, Hasenbein, Nortrud, Liu, Christine, Truong, Thy T., Galway, Moira E., Mansfield, Shawn D., Hocart, Charles H. & Wasteneys, Geoffrey O. The anisotropy1 D604N mutation in the Arabidopsis cellulose synthase1 catalytic domain reduces cell wall crystallinity and the velocity of cellulose synthase complexes. Plant Physiol. 162, 74–85 (2013).

64. Fujita, Miki, Himmelspach, Regina, Hocart, Charles H., Williamson, Richard E., Mansfield, Shawn D. & Wasteneys, Geoffrey O. Cortical microtubules optimize cell-wall crystallinity to drive unidirectional growth in Arabidopsis. Plant J. 66, 915–928 (2011).

65. Liu, Zengyu, Schneider, Rene, Kesten, Christopher, Zhang, Youjun Yi, Somssich, Marc, Zhang, Youjun Yi, Fernie, Alisdair R. & Persson, Staffan. Cellulose-Microtubule Uncoupling Proteins Prevent Lateral Displacement of Microtubules during Cellulose Synthesis in Arabidopsis. Dev. Cell 38, 305–315 (2016).

66. Standal, R., Iversen, T. G., Coucheron, D. H., Fjaervik, E., Blatny, J. M. & Valla, S. A new gene required for cellulose production and a gene encoding cellulolytic activity in Acetobacter xylinum are colocalized with the bcs operon. J. Bacteriol. 176, 665–672 (1994).

67. López, Daniel, Vlamakis, Hera, Losick, Richard & Kolter, Roberto. Cannibalism enhances biofilm development in bacillus subtilis. Mol. Microbiol. 74, 609–618 (2009).

68. Billings, Nicole, Birjiniuk, Alona, Samad, Tahoura S., Doyle, Patrick S. & Ribbeck, Katharina. Material properties of biofilms - A review of methods for understanding permeability and mechanics. Reports Prog. Phys. 78, (2015).

69. Stewart, Philip S. Diffusion in biofilms. Journal of Bacteriology vol. 185 1485–1491 (2003).

70. Xu, Karen D., Stewart, Philip S., Xia, Fuhu, Huang, Ching Tsan & McFeters, Gordon A. Spatial physiological heterogeneity in Pseudomonas aeruginosa biofilm is determined by oxygen availability. Appl. Environ. Microbiol. 64, 4035–4039 (1998).

71. Jo, Jeanyoung, Cortez, Krista L., Cornell, William Cole, Price-Whelan, Alexa & Dietrich, Lars E. P. An orphan cbb3-type cytochrome oxidase subunit supports Pseudomonas aeruginosa biofilm growth and virulence. Elife 6, (2017).

72. Sartori, N., Richter, Karsten & Dubochet, Jacques. Vitrification depth can be increased more than 10-fold by high-pressure freezing. J. Microsc. 172, 55–61 (1993).

73. Harapin, Jan, Börmel, Mandy, Sapra, K. Tanuj, Brunner, Damian, Kaech, Andres & Medalia, Ohad. Structural analysis of multicellular organisms with cryo-electron tomography. Nat. Methods (2015).

74. Hsieh, Chyongere, Schmelzer, Thomas, Kishchenko, Gregory, Wagenknecht, Terence & Marko, Michael. Practical workflow for cryo focused-ion-beam milling of tissues and cells for cryo-TEM tomography. J. Struct. Biol. 185, 32–41 (2014).

75. Schaffer, Miroslava, Pfeffer, Stefan, Mahamid, Julia, Kleindiek, Stephan, Laugks, Tim, Albert, Sahradha, Engel, Benjamin D., Rummel, Andreas, Smith, Andrew J., Baumeister, Wolfgang & Plitzko, Juergen M. A cryo-FIB lift-out technique enables molecular-resolution cryo-ET within native Caenorhabditis elegans tissue. Nat. Methods 16, 757–762 (2019).

76. Das, Aditi & Das, Anath. Delineation of polar localization domains of *Agrobacterium tumefaciens* type IV secretion apparatus proteins VirB4 and VirB11. Microbiologyopen 3, 793–802 (2014).

77. Rigort, Alexander, Bäuerlein, Felix J. B., Leis, Andrew, Gruska, Manuela, Hoffmann, Christian, Laugks, Tim, Böhm, Ulrike, Eibauer, Matthias, Gnaegi, Helmut, Baumeister, Wolfgang & Plitzko, Jürgen M. Micromachining tools and correlative approaches for cellular cryo-electron tomography. J. Struct. Biol. 172, 169–179 (2010).

78. Danev, Radostin, Buijsse, Bart, Khoshouei, Maryam, Plitzko, Jürgen M. & Baumeister, Wolfgang. Volta potential phase plate for in-focus phase contrast transmission electron microscopy. Proc. Natl. Acad. Sci. U. S. A. 111, 15635–40 (2014).

79. Mastronarde, David N. Automated electron microscope tomography using robust prediction of specimen movements. J. Struct. Biol. 152, 36–51 (2005).

80. Kremer, J. R., Mastronarde, D. N. & McIntosh, J. R. Computer visualization of three-dimensional image data using IMOD. J. Struct. Biol. 116, 71–6 (1996).

