## Supplemental figures for "Structure of the bacterial cellulose ribbon and its assembly-guiding cytoskeleton by electron cryotomography"

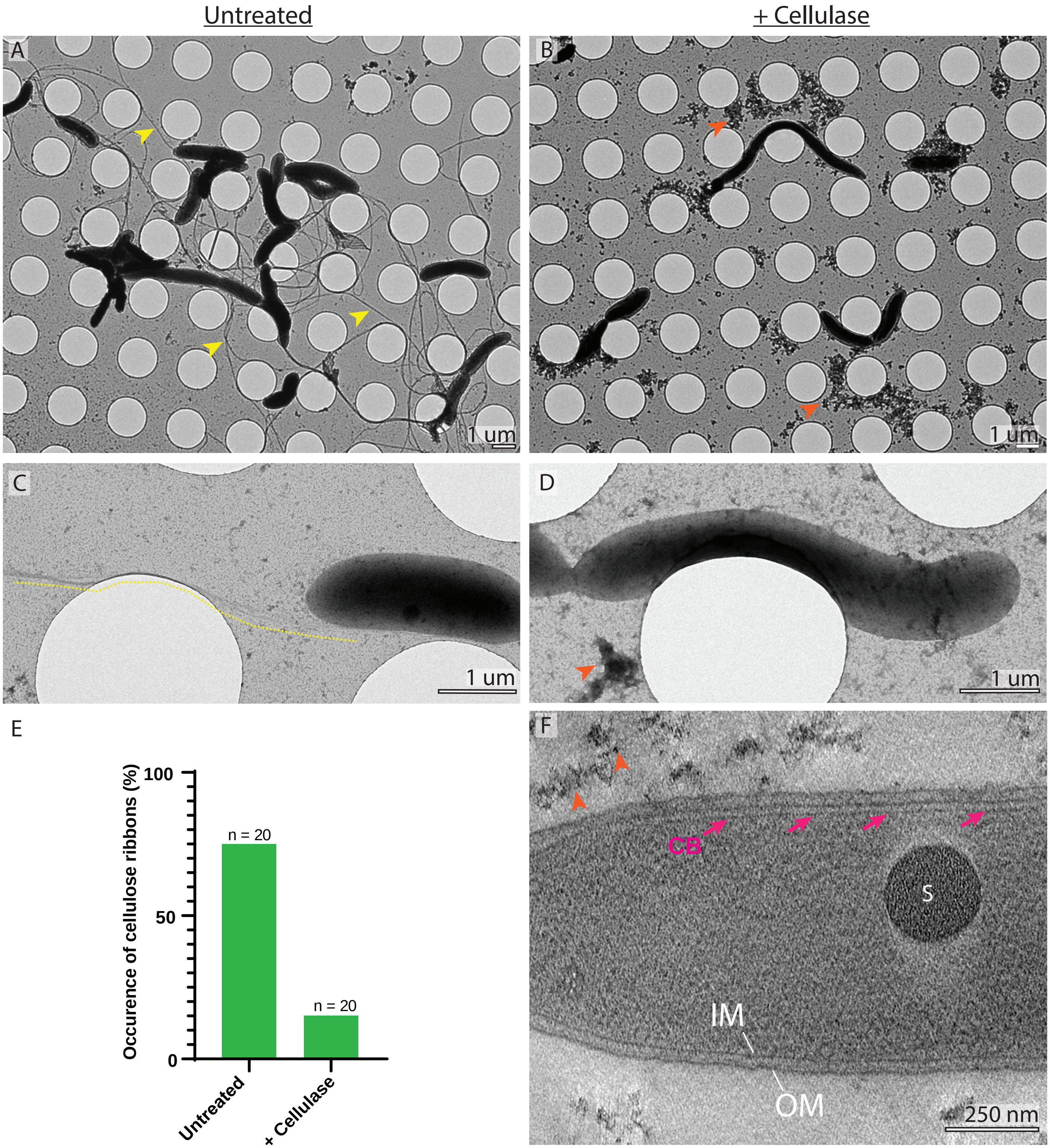
Untreated
+ Cellulase
A
B
1 um
1 um
C
D
1 um
1 um
E
Occurence of cellulose ribbons (%)
100
50
0
n = 20
n = 20
Untreated
+ Cellulase
F
CB
S
IM
OM
250 nm

**Supplemental figure 1 | Cellulase treatments prevents the aggregation of cells and the occurrence of the cellulose ribbons along the side of the cells**

(**A**) Low mag image of a negatively stained *G. hansenii* culture 300 minutes post-separation. Dense fibers visible alongside the cells (yellow arrowheads) hold the cells together. (**B**) Low mag image of a negatively stained *G. hansenii* culture 300 minutes post-separation treated with 0.2g/L cellulase. No fibers are visible but putative clusters of degraded cellulose are seen (orange arrowheads). (**C**) Enlarged view of an untreated cell displaying a cellulose ribbon on its side (yellow dashed line underlining the cellulose ribbon). (**D**) Enlarged view of a cell after cellulase treatment where the clusters of degraded cellulose are seen (orange arrowhead). (**E**) Occurrence of cellulose in 20 overviews like (A) and (B) for each condition. (**F**) 11-nm thick cryo-tomographic slice of a *G. hansenii* cell treated with 0.2g/L cellulase exhibiting putative degraded cellulose aggregates (orange arrowheads), already observed in previous studies where *G. xylinum* cellulose is digested by cellulase[47], and the cortical belt (purple arrows).

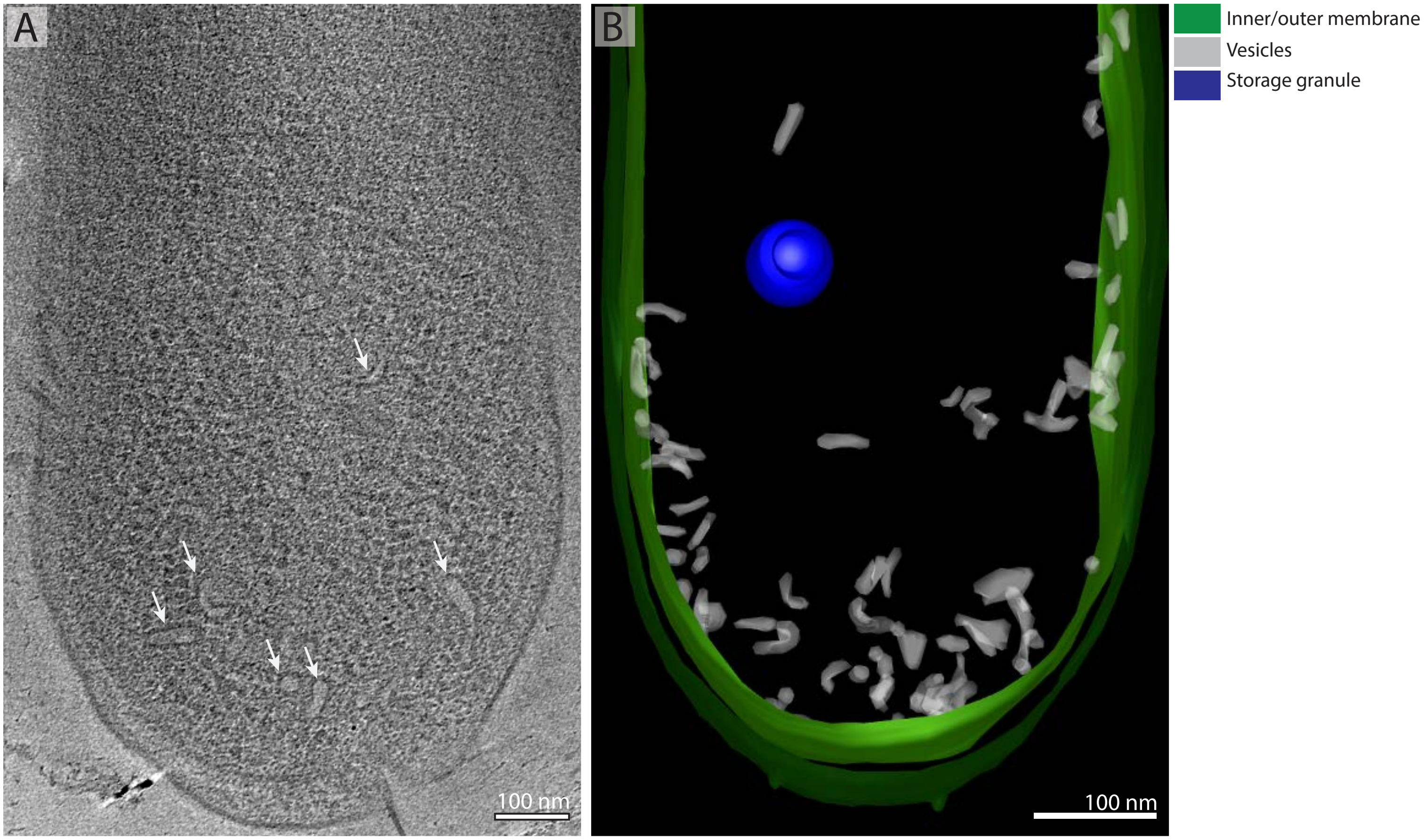


**Supplemental figure 2 | *Gluconacetobacter hansenii* cells harbor cytoplasmic vesicles**

(**A**) 11- nm thick tomographic slice of a *G. hansenii* cell showing numerous vesicles in the cytoplasm (white arrows). (**B**) Manual 3D segmentation of the tomogram shown in (A).

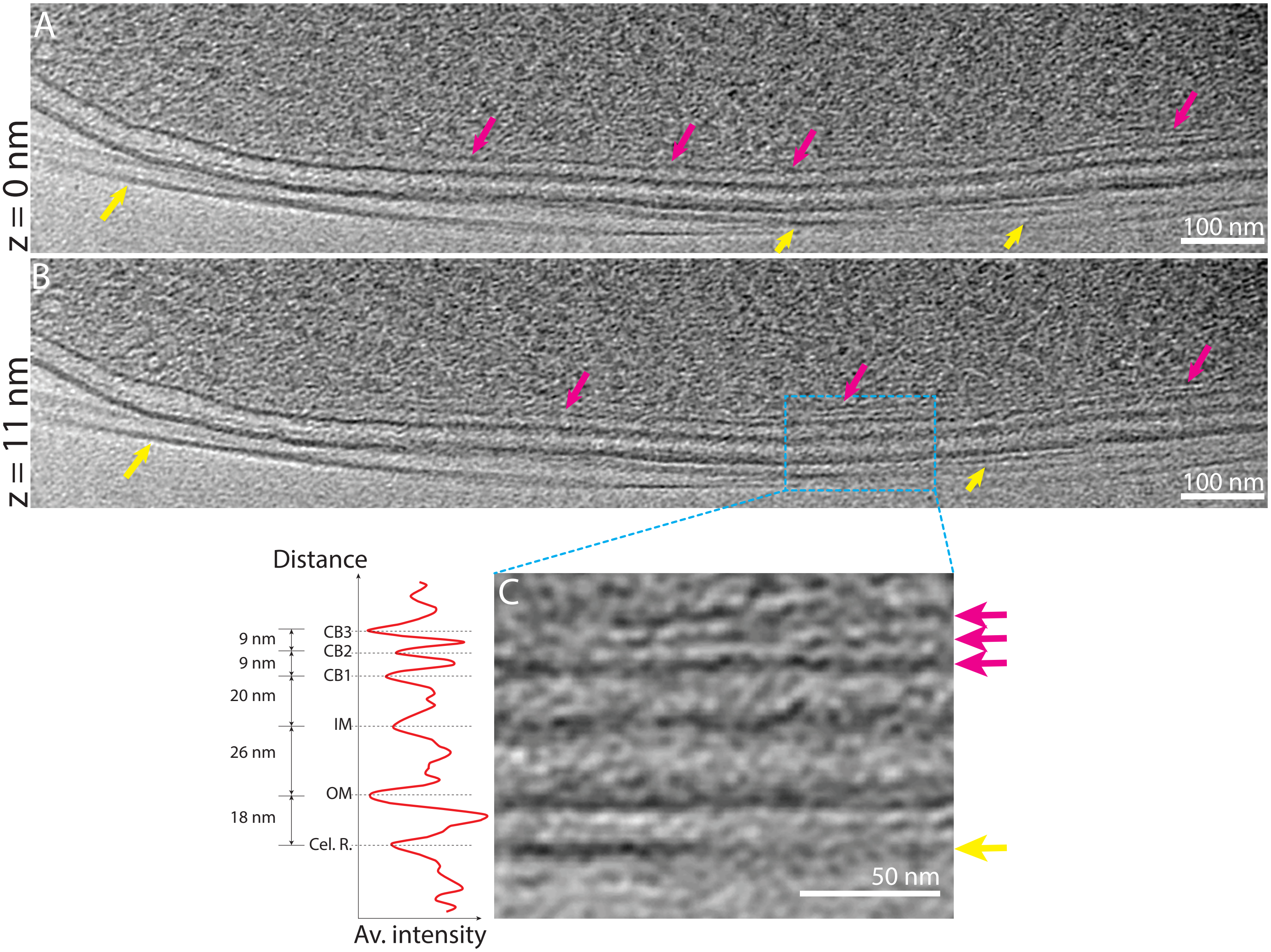


**Supplemental figure 3 | *Gluconacetobacter xylinus* has the same layout of the cellulose and the cortical belt as of *G. hansenii*.**

(**A-B**) Two 11-nm thick tomographic slices of the same *G. xylinus* cell. The cellulose sheet and the cortical belt are indicated by yellow and purple arrows, respectively. (**C**) Enlarged view of the blue boxed region in (B) showing the layering of the cortical belt and the juxtaposition of the cortical belt and the cellulose ribbon. On the left is the density profile.
